# GRASP: A PLANT TRANSFORMATION-INDEPENDENT CRISPR-BASED SYSTEM FOR AFFINITY PURIFICATION OF SPECIFIC CHROMATIN LOCI

**DOI:** 10.64898/2026.03.18.712347

**Authors:** Aurélien Devillars, Silvia Farinati, Adriana Fernanda Soria Garcia, Justin Joseph, Giovanni Gabelli, Sara Zenoni, Edoardo Bertini, Alessandra Amato, Bhanu Prakash Potlapalli, Andreas Houben, Fabio Palumbo, Gianni Barcaccia, Alessandro Vannozzi

## Abstract

Chromatin organization regulates genome stability and gene expression by controlling DNA accessibility to transcription factors and regulatory complexes. DNA-protein interactions are commonly investigated using chromatin immunoprecipitation (ChIP), which relies on specific antibodies often involving technically demanding protocols. CRISPR-Cas technologies have enabled sequence-specific targeting of genomic loci using catalytically inactive Cas9 (dCas9), but most CRISPR-based chromatin capture approaches in plants require transient or stable transformation to express the CRISPR machinery, limiting their applicability across species, tissues and physiological contexts. Here, we present GRASP (Genomic Region Affinity Sequestration by CRISPR-Purification), a transformation-independent strategy for sequence-specific chromatin isolation operating directly on purified plant nuclei. In GRASP, dCas9-gRNA ribonucleoprotein complexes are used to capture predefined genomic regions from chromatin under native conditions, bypassing the need for transgene expression. Using grapevine and tomato as model systems, we demonstrate efficient and highly specific enrichment of target loci, including telomeric repeats as well as low-copy and single-copy genomic regions, with qPCR and NGS validation. These results establish GRASP as a robust and broadly applicable platform for locus-specific chromatin isolation in plants. Beyond sequence-specific DNA isolation, GRASP establishes a versatile platform for potential downstream analyses of locus-associated chromatin components, including protein complexes, distal DNA-DNA interactions and chromatin-associated RNAs, providing new opportunities to investigate regulatory architecture in plant genomes.

**GRAPHICAL ABSTRACT:** 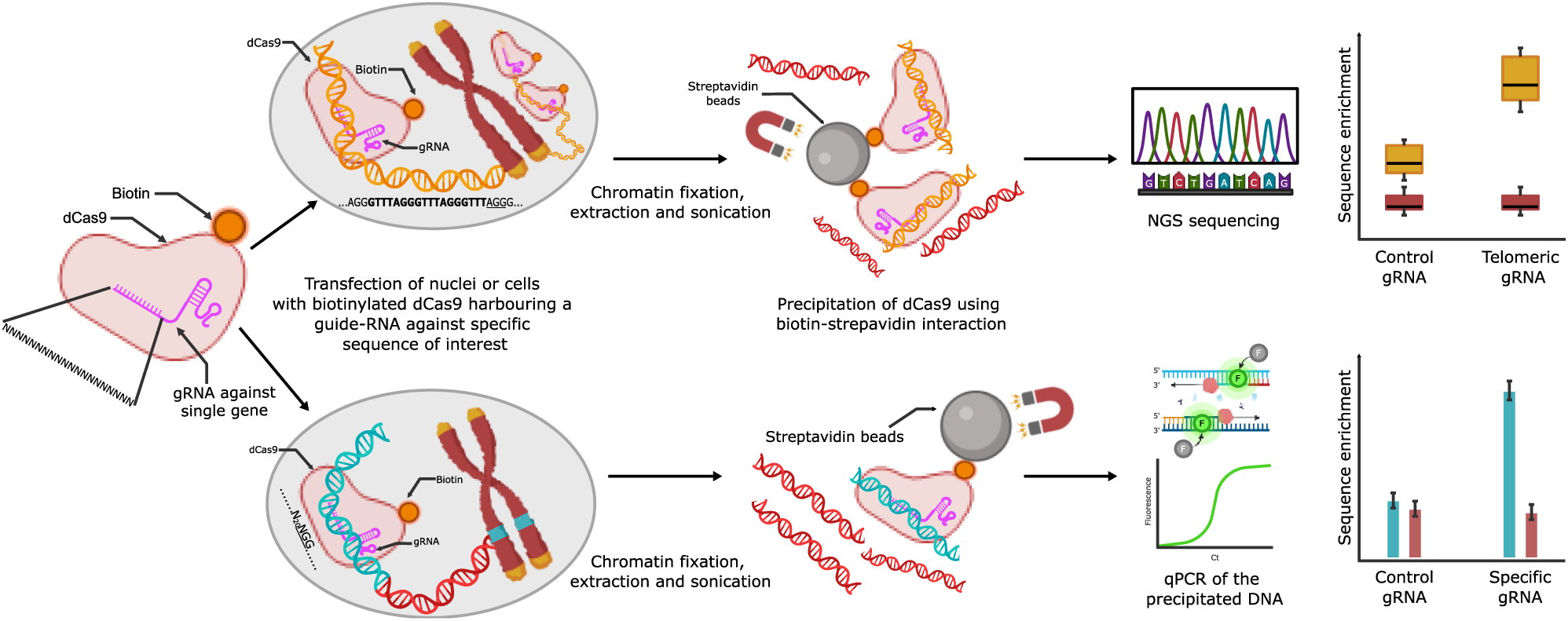

## INTRODUCTION

Chromatin is a complex, three-dimensional and dynamic structure composed of DNA, RNA, and proteins organized to regulate genome compaction through mechanisms that are broadly conserved across eukaryotes. This organization plays a key role not only in DNA replication, repair and recombination, but also in gene expression, gene regulation and the special organization of the genome itself (1). Rather than occurring through independent pathways, these processes are now understood as taking place in a functionally and physically connected manner, collectively contributing to the regulation of gene expression and the maintenance of nuclear homeostasis (2).

With respect to gene expression, a major determinant of gene regulation is the accessibility of genomic loci to transcription factors and the transcriptional machinery. Transcriptionally active regions are generally associated with relatively less compact chromatin (euchromatin), whereas more compact chromatin regions (heterochromatin) are largely transcriptionally inactive and can account for a substantial fraction of the genome in species with large genomes (2). The chromatin state of a given sequence largely depends on the nature of the proteins and complexes present at that locus, as well as on the post-translational modifications carried by histones (3). Beyond transcriptional regulation, chromatin organization also play a key role in genome stability, for example at the level of telomeres, where specific protein complexes promote a specialized organization and compaction of chromosome ends, protecting them from degradation (4, 5). The classical approaches used to investigate DNA‒protein interactions, and thus, chromatin state and gene regulation, include electromobility mobility shift assays (EMSAs), DNase footprinting techniques, reporter gene-based assays and chromatin immunoprecipitation (ChIP), the latter being among the most widely used methods (6). ChIP relies on specific antibodies to recognize a protein of interest and precipitate the chromatin fragment to which it is bound. When coupled to next-generation sequencing (ChIP-seq), this approach enables genome-wide identification of protein binding sites. Despite its versatility and broad applicability across biological systems, the method presents several practical limitations, including the technical complexity of the protocol and its reliance on the availability and specificity of suitable, which can hinder its effective implementation (7).

Over the past decade, in addition to genome editing, CRISPR-Cas systems have been repurposed to target nucleic acids in a sequence-specific manner for a wide-range of applications, including DNA and RNA imaging, modulation of gene activity *in vivo*, and depletion of undesired sequences from sequencing libraries (8, 9). Chromatin capture approaches have also emerged as part of these new generation tools, with several techniques developed to target and isolate specific genomic regions using a non-cutting version of Cas9, commonly referred to as “dead Cas9” (dCas9). These approaches have been implemented primarily in mammalian systems, but also in plant species (10–12). dCas9 is a catalytically inactive variant of the Cas9 protein that lacks endonuclease activity due to specific mutations in its nuclease domains (RuvC1 and HNH, typically D10A and H841A). Unlike wild-type Cas9, which introduce double-strand brakes at the target sites, dCas9 retains the ability to bind DNA in a sequence-specific manner through guidance by a single guide RNA (gRNA) (13).

Inspired by previous studies, we aimed to develop a protocol for chromatin isolation leveraging a dCas9-based system applicable in plant samples, focusing on two species of major agronomic relevance, namely grapevine (*Vitis vinifera* L.) and tomato (*Solanum lycopersicum* L.). Previous approaches for chromatin precipitation in plants have predominantly relied on stable, or more frequently transient, transformation-based approaches, in which the dCas9 machinery is delivered through expression vectors. Although effective, these methods are inherently limited by the requirement of efficient transfection protocols, which are not available for all plant species and tissues. Additional limitations arise when these approaches are used to study transcription factors (TFs) binding to *cis*-regulatory elements (CREs) of target genes. Since TF binding often depends on gene induction and many genes are activated only in specific tissues, organs, or environmental conditions, it is not always feasible to achieve transformation under the precise biological contexts in which these interactions occur. Moreover, transient transformation itself can introduce artificial alterations in gene expression, further complicating the interpretation of results (14, 15). To overcome these limitations and enhance the transferability of the technique across plant systems, we developed GRASP (Genomic Region Affinity Sequestration by CRISPR-Purification), a protocol for locus-specific chromatin isolation using purified nuclei as the starting material. By bypassing the need for transformation, this approach preserves the native chromatin environment and avoids potential transcriptional perturbations associated with transient expression systems. As a result, the method provides a framework for studying protein–DNA interactions in their physiological and developmental context and can be applied to species and tissues where transient transformation remains inefficient or where gene induction conditions are difficult to reproduce *ex vivo*.

As proof of concept, we initially targeted telomeric arrays, which represent highly repetitive genomic regions that are well suited for testing sequence-specific chromatin capture approaches. Telomeres consist of tandem repeats of a short sequence motif present in a tissue- and cell-type–dependent number of copies: TTAGGG_n_ in most vertebrates and TTTAGGG_n_ in most plants. Furthermore, telomeric chromatin typically displays an intermediate compaction state between euchromatin and heterochromatin (4, 16). Telomeric repeats have also been successfully visualized in living and fixed plant nuclei using CRISPR–dCas9 systems and have already been chosen as model targets for the development of CRISPR-Cas-system based methodologies (11, 17–22). To summarize our approach, we first performed a bioinformatics analysis to estimate telomere lengths in the species under investigation. We then validated the ability of dCas9, loaded with a specific gRNA spacer previously described in literature, to bind to telomeric sequences (20), first *in vitro*, by targeting a PCR-engineered DNA fragment and subsequently *in situ* on fixed nuclei by CRISPR–FISH (originally named RGEN-ISL (18)), an effective method for visualizing specific DNA regions in plant cells (18, 20).

After establishing the system on telomeric repeats, we extended the approach to more complex genomic targets, including low-copy and single-copy loci. We then developed an affinity-based precipitation protocol for sonicated plant chromatin and validated its specificity by targeting these different genomic regions. Enrichment of the target sequences in the precipitated chromatin fractions was assessed through next-generation sequencing (NGS) or qPCR analyses of the recovered DNA. Together, these results represent an important step toward the development of efficient sequence-specific chromatin isolation methodologies in plants, with potential applications for investigating genome organization and gene regulatory mechanisms.

## METHODS

### Reagents and solutions

All reagents, kits and solutions, including their abbreviations, used in this study are detailed in **Supplementary Methods 1**, **2** and **3**, respectively. Several parts of the protocols are also reported in **Supplementary Method 4** instead of the main manuscript.

### Plant material

Leaf samples were collected from explants of fully grown *in vitro* plants. The plants were grown on Murashige and Skoog medium (4.4 g/L MS with vitamins), supplemented with sucrose (20 g/L), IBA (0.1 mg/L), and cefotaxime (100 mg/L) and solidified with 7 g/L plant agar. Cultures were grown in a controlled growth chamber at 25 °C under a 12 h light/12 h dark photoperiod, with a light intensity of 35 µmol.m^-2^s^-1^.

### Counting of the telomeric repeats in grapevine T2T genomes

We developed an *in silico* pipeline to quantify the number telomeric motif repeats in each genome and to map their distribution along the chromosomes. To avoid counting isolated or spurious occurrences, we focused on 21-mers corresponding to three consecutive repetitions of the telomeric motif “TTAGGG”. Depending on the starting position within the repeat unit, this results in 14 distinct telomeric 21-mers: TTTAGGGTTTAGGGTTTAGGG, TTAGGGTTTAGGGTTTAGGGT, TAGGGTTTAGGGTTTAGGGTT, AGGGTTTAGGGTTTAGGGTTT, GGGTTTAGGGTTTAGGGTTTA, GGTTTAGGGTTTAGGGTTTAG, GTTTAGGGTTTAGGGTTTAGG, and their reverse-complement equivalents. The main limitation of this pipeline is that it requires a telomere-to-telomere (T2T) genome assembly, i.e. a fully contiguous sequence spanning the entire length of each chromosome. The pipeline was implemented as follows. First, we used Jellyfish (version 2.3.0) to count the occurrences of all 21-mer in the genomes of *V. vinifera* cv. Pinot Noir (23), cv. Thompson Seedless (24), *Arabidopsis thaliana* (25) and *Zea mays* inbred line Mo17 (26), retaining only those 21-mer occurring at least 1000 times. Next, telomeric 21-mers identified in the previous step were used as queries in BLAST searches against the corresponding genomes, requiring 100% sequence identity between the query and the matched 21-mer. The resulting BLAST hits were further filtered using a custom R script to remove (i) alignments shorter than 21 bp, which do not span the full length of the telomeric 21-mers, and (ii) overlapping hits, expected given the repetitive nature of sequences composed of three consecutive ‘TTAGGG’ motifs. The genome was then partitioned into fixed-size sectors, and for each sector of each chromosome, the number of telomeric 21-mers was quantified. A graphical representation of the chromosome-wide telomeric repeat distribution was subsequently generated in R using the ggplot2 package, with chromosomes displayed as sectorized tracks and associated repeat counts. In addition, the genomic coordinates of telomeric arrays were inferred by the R script and defined as the positions of the first and last 21-mers within the sectors adjacent to chromosome ends. The bash-based and R-based components of the pipeline are provided in **Supplementary File 1**.

*(Troubleshooting steps that may be required are mentioned in* **Supplementary Methods 5**)

### *In vitro* amplification of telomeric repeat-containing fragments

A detailed description of this procedure is provided in **Supplementary Method 4.**

### RNP preparation and complex assembly

- Use a two-part guide RNA (gRNA) system composed of a telomere-specific crRNA (GGGTTTAGGGTTTAGGGTTT) (18) and tracrRNA-ATTO550, along with biotinylated dCas9.
- Reconstitute lyophilized crRNA, tracrRNA and biotinylated dCas9 protein according to the manufacturers’ instructions.
- Mix equal volumes of 200 µM crRNA and 200 µM tracrRNA in a sterile PCR tube.
- Incubate at 95 °C for 5 min, then at room temperature (RT) for 20 min to form a 100 µM gRNA stock solution. Store the gRNA stock at −20 °C until use.
- To prepare the working solution, dilute 2 µL of the 100 µM stock with 18 µL of nuclease-free duplex buffer (provided by the manufacturer) to obtain a 10 µM working aliquot.
- To assemble the RNP complex, mix 1 µL of 10 µM gRNA, 1 µL of 6.25 µM dCas9 protein in 100 µL of RNP solution, in a 1.5 mL Low Protein Binding Microcentrifuge Tube.
- Incubate the mixture at 26 °C for 10 min then keep the assembled RNP complex on ice until use.
- When performing CRISPR-FISH, a volume of 100 µL of the RNP complex is sufficient for 4 slides.

### Cleavage assay with Cas9 and gRNA targeting telomeres

Following RNP assembly, we evaluated the cleavage activity of the Cas9:gRNA complex on a telomeric target *in vitro*. The detailed protocol for the cleavage assay using Cas9 and the telomere-targeting gRNA is provided in **Supplementary Method 4**. The sequences of all primers and DNA fragments used in this assay are listed in **Supplementary File 2**.

### Binding assay

A comprehensive description of the experimental procedure is available in **Supplementary Method 4**.

### Nuclei isolation from plant leaves for CRISPR–FISH and FISH

Nuclei were isolated from approximately 10 mg of healthy young leaves of tomato, *V. vinifera* (cv. Thompson Seedless, “Sultanna”). The protocol was adapted from Ishii et al. (2019) for use in grapevine and was performed as follows:

- Fix the leaves in ice-cold formaldehyde solution (2% for tomato or 4% for grapevine) diluted in Tris buffer. Apply vacuum infiltration for 5 min followed by continued fixation for 25 min on ice without vacuum.
- Rince the tissues twice for 5 min each in ice-cold Tris-buffer.
- Finely chop the tissues in 400–500 µL ice-cold LB01 buffer (Doležel et al. 1989) with a fresh blade.
- Filter the suspension through a 35 µm pore size mesh.
- Transfer 100 µL of the filtered nuclei suspension per slide onto glass microscope slides and centrifuge using a Cytospin 4 (Epredia, cat. no. A78300003) at 700 rpm for 5 min.
- Mark the position of the sample by raying the slide on both sides of the halo formed by the nuclei on the slide.
- Keep the slides in ice-cold PBS until use.

### CRISPR–FISH

CRISPR–FISH was performed as described previously by (27). A brief overview of the principle of this technique is provided in **Supplementary Method 4**.

### Fluorescence In Situ Hybridization (FISH)

FISH was performed as described in (18) with following modifications:

- Do not incubate the nuclei with pre-hybridization mixture.

- Prepare a telomere-specific, ATTO550-dUTP-labelled probe following the method described in (28).

### Colocalization of CRISPR–FISH and FISH signals

- Perform CRISPR-FISH on the desired species as described above. Using a motorized-stage fluorescence microscope, identify and record the coordinates of approximately 10 fields containing nuclei with clear and well-defined signals. Capture images of the DAPI and CRISPR-FISH channels for each selected field.
- Carefully remove the coverslip by washing the slides in PBS for 5 min at RT.
- Repeat the PBS wash once more under the same conditions.
- Proceed with FISH as previously described, starting from the fixation step in ethanol:acid (3:1).
- Using the saved coordinates, relocate the image of the same nuclei fields. Capture images of the DAPI and FISH channel.
- Align the CRISP-FISH and FISH based on DAPI channels as a reference. In our analysis, alignment was performed using GIMP (v2.10.34) and ImageJ/FIJI (v1.54f).
- A detailed step-by-step procedure for image alignment is described in **Supplementary Methods 3**.
- To qualitatively assess colocalization between CRISPR-FISH and FISH foci, use ImageJ to extract pixel intensity profiles from both channels along a defined line that should intersect multiple nuclear foci.
- Export the resulting values as a data table and plot intensity profiles using the ggplot2 package in R.

### Nuclei extraction from plant leaves for the nuclei transfection assay

The detailed protocol for this procedure is provided in **Supplementary Method 4**.

### Nuclei transfection assay

The detailed protocol for this procedure is provided in **Supplementary Method 4**.

### Microscopy

Microscopy images were acquired using a Zeiss AXIO Observer. A Z1/7 inverted fluorescence microscope equipped with a ZEISS Axiocam 705 mono camera and a Plan-Apochromat 40×/1.4 Oil DIC M27 objective. CRISPR–FISH signals (ATTO500) and standard FISH signals (ATTO488 or ATTO550) were analyzed using the appropriate fluorescence filters. All fluorescence images were acquired in grayscale mode and subsequently pseudo-colored using ImageJ for downstream analysis and visualization.

*(Troubleshooting steps that may be required are mentioned in* **Supplementary Methods 5**)

### Nuclei transfection and chromatin streptavidin isolation for the precipitation of telomeric sequences

#### Nuclei isolation and transfection using a CRISPR-FISH-like protocol

- Collect at least 500 mg of leaf tissue, ideally around 1 g. This quantity is sufficient to obtain enough nuclei for 2 transfections, including the negative control.
- Fix the material in 1% formaldehyde prepared in ice-cold Tris-Buffer for 5 min under vacuum, followed by 25 min on ice w/o vacuum.
- Wash the fixed leaves twice in ice-cold Tris-Buffer for 5 min each.
- Transfer part of the leaf sample into a precooled Petri dish and finely chop it in 200-300 µL of ice-cold LB01 Buffer with a fresh blade.
- Add up to 600 µL ice-cold LB01 buffer to resuspend the released nuclei.
- Filter the suspension through a 35 µm mesh filter using a cut-off pipette tip.
- Repeat the chopping and filtering steps as needed until all leaf material is processed, changing the filter and the collection tube for each batch.
- Centrifuge the filtrate at 3000 × g for 10 min at 4°C.
- Resuspend the pellet in 500 µL ice-cold PBS. Pool nuclei from all batches, then split the suspension into two equal aliquots in 2 mL microcentrifuge tubes.
*(Troubleshooting steps that may be required are mentioned in* **Supplementary Methods 5**)
- Centrifuge at 3000 × g for 2 min at 4 °C.
- Prepare two aliquots of 400 µL block solution and two aliquots of 100 µL RNP solution (compositions specified in **Supplementary Method 4**), for the telomere targeting RNP complex, add 4 µL of 3xFLAG-biotinylated dCas9 and 4 µL of the assembled telomere-specific gRNA to the RNP solution. For the negative control RNP complex, add 4 µL of the 3xFLAG-biotinylated dCas9 and 4 µL of the assembled gRNA targeting *GAL4*.
- Incubate both RNP solutions at 26°C for 10 min, then place on ice until use.
- Resuspend each nuclei solution in 400 µL of block solution and incubate for 2 min at RT.
- Add 100 µL of the telomere-targeting RNP complex to one nuclei aliquot (sample precipitated with gRNA). Add 100 µL of the *GAL4*-targeting RNP complex to the second nuclei aliquot (negative controls sample, precipitated without gRNA).
- Incubate the samples at 37°C for 1-2 hours on a rotating wheel.
*From this point onward, process each nuclei aliquot separately*.
- Add 13.88 μL of 37% formaldehyde (final concentration: 1%) and invert on the wheel for 15 min, RT.
- Add 102.8 µL of 600 mM glycine (final concentration: 0.1 M) and invert on the wheel for 5 min at RT.
- Centrifuge at 3000 × g for 2 min.
- Resuspend the pellet in 500 µL PBS

#### Chromatin extraction from purified and transfected nuclei

- Centrifuge at 3000 × g for 5 min.
- Resuspend the pellet in 1 mL nuclei wash buffer with Triton.
- Incubate at 37°C for 30 min.
- Centrifuge at 3000 × g for 5 min.
- Resuspend the pellet in 1 mL nuclei wash buffer without Triton. Mix immediately by gently tapping the tube.
- Centrifuge at 3000 × g for 2 min.
- Resuspend the pellet in 300 µL nuclear lysis buffer. Pipette up and down.

#### Sonication

- Transfer the chromatin samples into tubes suited for sonication
- Sonicate on a Bioruptor*®* Plus at high intensity for 35 cycles (30 sec ON; 30 sec OFF). *(Troubleshooting steps that may be required are mentioned in* **Supplementary Methods 5**)
- Briefly centrifuge samples before opening.
- Add 100 µL nuclear lysis buffer and transfer to a new, standard microcentrifuge tube. The samples may be frozen at −80°C until use.
- Add 44 µL of 10% Triton X-100 to sonicated nuclei lysate and centrifuge 10 min at 16,100 × g at 4 °C.
- Carefully transfer 400 µL of the supernatant into a new microcentrifuge tube and add 24 µL of 5M NaCl; this is the soluble chromatin.
- Take 20 µL residual supernatant as the input control and store at −80 °C in a 0.5 mL tube. *(Troubleshooting steps that may be required are mentioned in* **Supplementary Methods 5**)

### Streptavidin-mediated precipitation of telomeric sequences

- For each sample, wash 12.50 µL Dynabeads™ MyOne™ T1 magnetic beads in 2mL Low Protein Binding Microcentrifuge Tubes microcentrifuge tubes as follows:

- Resuspend the beads in the vial (vortex for >30sec, or tilt and rotate for 5 min).
- Transfer the desired bead volume to a tube.
- Resuspend in 1 mL of RIPA 0.3 buffer and vortex.
- Place the tube on a magnet for 1 min and discard the supernatant.
- Resuspend again in 1 mL RIPA 0.3 buffer and vortex.
- Leave beads on the magnet in the RIPA 0.3 buffer from the second wash until use (at least 1 min). *(Troubleshooting steps that may be required are mentioned in* **Supplementary Methods 5**)
- After saving the input fraction from the sample, discard the RIPA 0.3 supernatant from the beads and remove the tubes from the magnet.
- Add the soluble chromatin directly to the beads and incubate tubes on a rotator at 4°C for at least 3 hours, or overnight.
- Collect beads on a magnet stand and discard supernatant.
- Wash beads by adding 1 mL of 2% SDS, vortex at maximum speed 15 sec and collect beads on a magnet. Discard the supernatant.
- Repeat precedent step as follows:

- One additional wash with 1 mL of 2% SDS
- Two washes with 1 mL High salt wash buffer.
- Two washes with 1 mL LiCl wash buffer.
- Two washes with 1 mL TE buffer. *(Troubleshooting steps that may be required are mentioned in* **Supplementary Methods 5**)
- Resuspend beads in 100 µL SDS elution buffer and incubate at 65 °C overnight with shaking.
- Briefly spin down to collect the beads. Separate the beads on magnet for 3 min and transfer supernatant to a new Low Protein Binding Microcentrifuge Tubes microcentrifuge tube.
- Rinse beads with 50 µL SDS elution buffer and pool supernatant with the eluted chromatin
- Add 1 µL of 0.5 μg/µL RNase A and incubate at 37 °C for 30 min.
- Add 2 µL of 10 mg/mL Protease K and incubate at 42 °C for 2 hours.
- Purify ChIP DNA using MinElute PCR Purification kit and elute DNA in 20 µL ddH_2_O.
- For input fraction reverse-crosslinking, add 1.62 µL of 2M NaCl (final concentration: 150 mM) to the thawed input and incubate at 65 °C overnight.
- Add an equal volume of phenol:chloroform:isoamyl alcohol (25:24:1), vortex and centrifuge for 10 min.
- Recover the upper phase. It is possible to repeat these two preceding steps to remove more of the protein.
- Add 2.5 volumes of 100% EtOH and 0.2 µL glycogen, then precipitate overnight at −20 or - 80 °C.
- Centrifuge at 4 °C for ≥30 min and discard supernatant.
- Wash the pellet with 500 µL of 70% EtOH and centrifuge for 15 min at 4 °C.
- Air-dry pellet at RT and resuspend DNA in 20 µL ddH_2_O.
- Note: DNA quantification is often unnecessary, as library preparations and sequencing are successful even when Qubit HS assay yields undetectable values. Store the rest of the sample at −20°C.
- Sequence DNA using WGS or ChIP-seq-compatible library preparation protocols and NGS. In this study, we used an Illumina platform that produced paired end reads.

### Bioinformatic analysis of sequencing data for telomeric sequence enrichment

- Trim raw sequencing reads using fastp software (version 0.20.0) with the following parameters:

~~~
--qualified_quality_phred 20, --unqualified_percent_limit 30, -- average_qual 25, --low_complexity_filter, --complexity_threshold 30, --length_required 40, --overrepresentation_analysis, and --detect_adapter_for_pe.
~~~

Use 10 threads to optimize the processing speed.

- For each sample, calculate 21-mer occurrence using Jellyfish (v 2.3.0)
- Retain only the 21-mers that repeated ≥ 1000 times.
- Process 21-mer counts using R as follows:
- Treat paired reads as technical replicates and calculate the mean occurrence values for each 21-mer.
- For each sample, normalize all occurrence values by dividing them by the mean occurrence value within the sample.
- Generate boxplot graphics the ggplot2 R package, categorizing the 21-mers into two groups: telomeric and non-telomeric 21-mers.

### Precipitation of low-copy spacers in grapevine

Indications for the design of spacers and primers, as well as sequences can be found in **Supplementary Method 4** and **Supplementary File 2** respectively.

- Perform precipitation of the target sequences using the different designed and synthesized gRNAs as described above, with following minor modifications:

- Split the isolated nuclei in as much aliquot of 10^8^ nuclei as used gRNA, counting them with a Denovix CellDrop counter. Increase the starting material if needed.
- Centrifuge at 3000 × g for 2 min at 4 °C.
- Prepare 400 µL block solution and 100 µL RNP solution for each different gRNA.
- Assemble the targeting RNP complexes by adding, for each, 4 µL of 3xFLAG-biotinylated dCas9 and 4 µL of 10 µM gRNA to one aliquot of 100 µL RNP solution.
- Incubate at 26°C for 10 min.
- Resuspend each aliquot of nuclei in 400 µL Block solution.
- Proceed with transfection and subsequent precipitation in the exact same way as performed with telomeres- and *GAL4*-specific bipartite spacers. Finaly elute precipitates and inputs in 50 µL ddH_2_O instead of 20 µL.
- After final elution, analyze all input and precipitate samples with qPCR using PowerUp™ SYBR™ Green Master Mix for qPCR using, for each well, 5 µL PowerUp mix, 0.6 µL Fw and 0.6 Rv primer, 1 µL of sample and ddH_2_O up to 10 µL. Here, the analyzes were set up, run and viewed on QuantStudio™ 3 Real-Time PCR System (machine and software). If the plate does not contain enough wells to allow for all samples and targets, perform qPCRs on precipitated and input samples in different plates.

*(Troubleshooting steps that may be required are mentioned in* **Supplementary Methods 5**)

## RESULTS

### Quantification and chromosomal distribution of telomeric repeats

The number of telomeric repeats in plants can vary greatly, not only across different species and varieties (29) but also among individuals and chromosomes within the same genome (25, 30–32). Moreover, variations in telomere length have been reported between different organs (33) and even between distinct cell types (34). Because the efficiency of dCas9-based chromatin capture is expected to depend, at least in part, on the density of available binding sites, we first assessed the abundance of telomeric repeats in grapevine. This analysis was intended to verify that grapevine telomeres, unlike those of tomato, which have already been successfully targeted in CRISPR–Cas applications, are present in sufficient copy numbers to support multiple dCas9 binding events and thus serve as a suitable proof-of-concept target for chromatin precipitation. We thus computed the number of all three-time repetitions of the Arabidopsis-type telomeric motif TTAGGG in the telomere-to-telomere genome assemblies of several plant species and genotypes, including the *V. vinifera* genotypes Sultana/Thompson Seedless (24) and PN40024 (23). In addition to the V. vinifera varieties, we included *A. thaliana* (35) and *Z. mays* (26). as reference species representing plant genomes with different sizes and telomere architectures. This comparative analysis provided a broader context for evaluating the abundance of telomeric repeat targets in grapevine and for estimating the potential density of dCas9 binding sites available for chromatin precipitation. The initial count of the telomeric 21-mers with the Jellyfish module followed by a BLAST validation of the identified 21-mers against the complete *V. vinifera* genome, revealed that telomeric-like sequences are more abundant in Thompson Seedless variety than in that of PN40024 line, with average occurrences of 24,000 and 15,800, respectively. We mapped the repetitions onto the chromosomes, allowing us to distinguish the telomeric arrays located at the chromosome ends from those occurring in internal regions. This approach provided a more accurate prediction of telomere length. Notably, we detected a higher abundance of telomeric sequences in Thompson Seedless (38 telomeric arrays over 16 Kbp in length) (**Figure 1**, **Table 1**) than in Pinot Noir (38 telomeric arrays over 13 Kbp in length on average) (**Figure S1**, **Table S1**).

**Figure 1.**
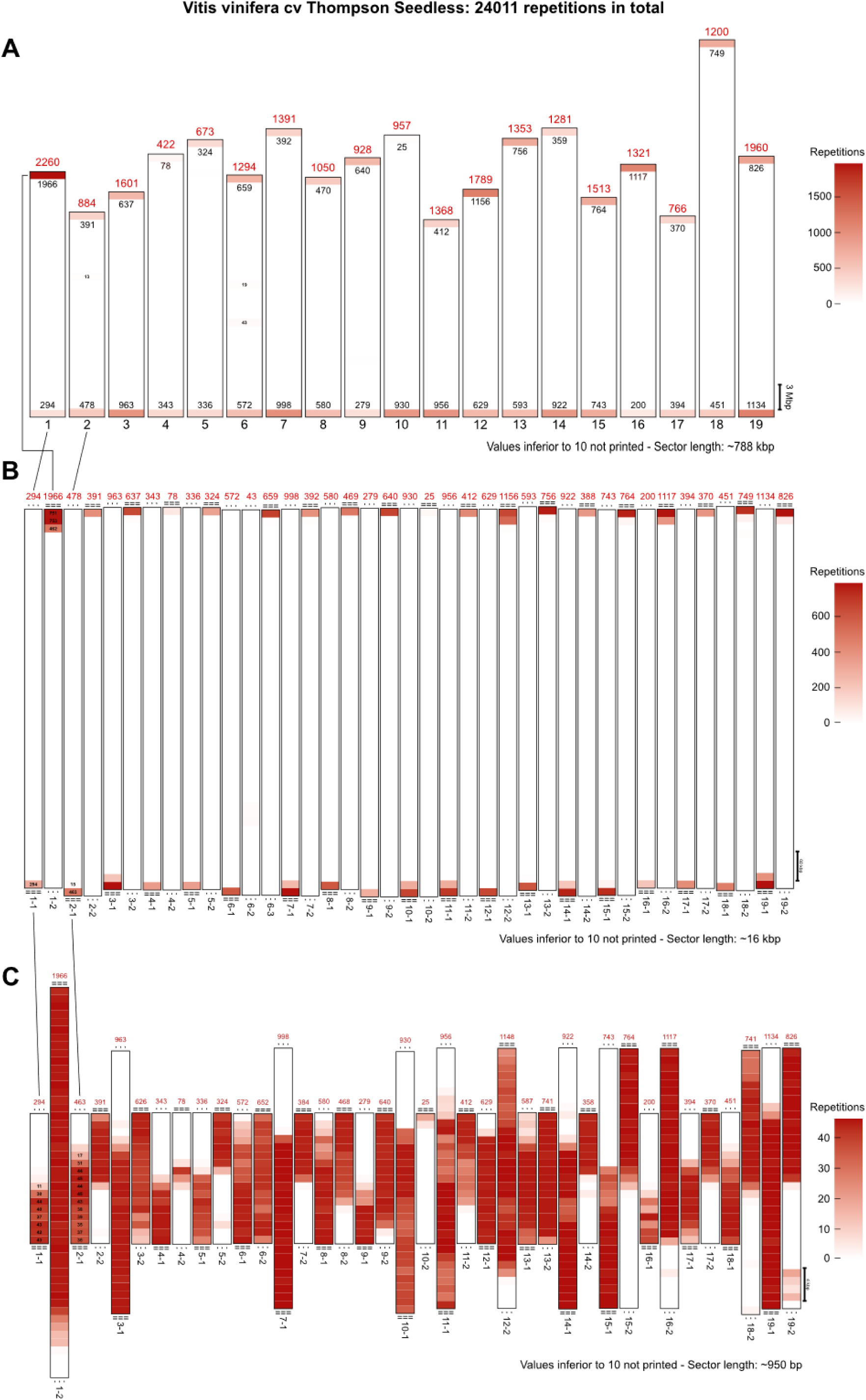
Abundance of telomeric repeats in the T2T genomes of *V. vinifera* cv Thompson Seedless. Occurrences of the telomeric 21-mer (AAACCCTAAACCCTAAACCCT) were identified by BLAST searches against the genome and mapped along chromosomes by partitioning each chromosome into fixed-size sectors (**A**). The number of 21-mer repeats within each sector is represented by a color scale (white to red), with higher intensity indicating higher repeat density. Sectors containing ≥20 repeats were further magnified (**B)**, and a higher-resolution view was used to highlight regions with persistent high repeat density (**C**). Red numbers indicate the total number of telomeric 21-mer repeats per chromosome or sub-chromosomal region, whereas numbers within each sector represent the local repeat counts. Ellipses (“···”) denote continuation of the chromosome beyond the displayed region, while “===” indicates that the chromosome starts or ends within the shown segment.

**Table 1.**
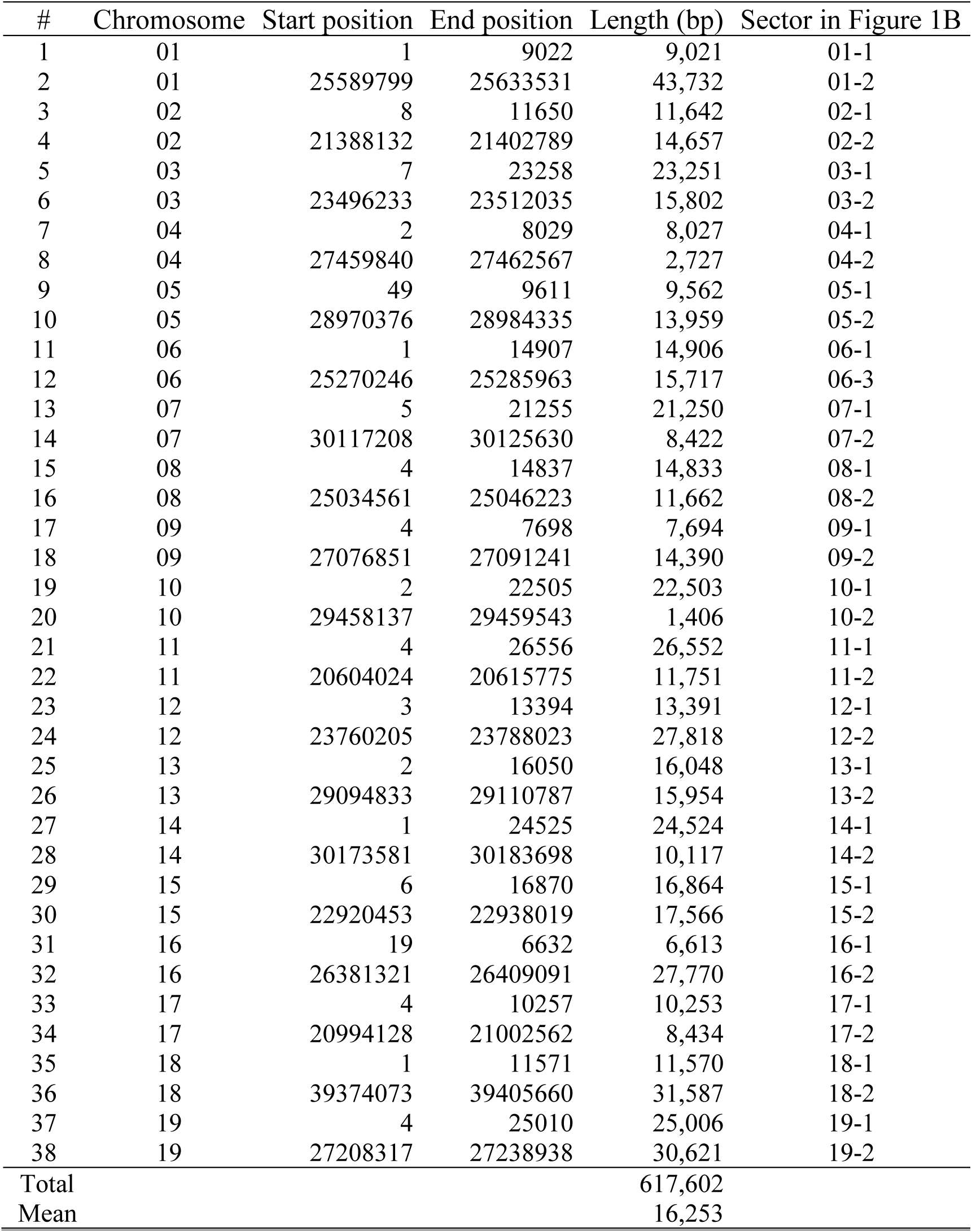
Telomeric regions in the *V. vinifera* cv. Thompson Seedless T2T genome assembly.

| # | Chromosome | Start position | End position | Length (bp) | Sector in Figure 1B |
| --- | --- | --- | --- | --- | --- |
| 1 | 01 | 1 | 9022 | 9,021 | 01-1 |
| 2 | 01 | 25589799 | 25633531 | 43,732 | 01-2 |
| 3 | 02 | 8 | 11650 | 11,642 | 02-1 |
| 4 | 02 | 21388132 | 21402789 | 14,657 | 02-2 |
| 5 | 03 | 7 | 23258 | 23,251 | 03-1 |
| 6 | 03 | 23496233 | 23512035 | 15,802 | 03-2 |
| 7 | 04 | 2 | 8029 | 8,027 | 04-1 |
| 8 | 04 | 27459840 | 27462567 | 2,727 | 04-2 |
| 9 | 05 | 49 | 9611 | 9,562 | 05-1 |
| 10 | 05 | 28970376 | 28984335 | 13,959 | 05-2 |
| 11 | 06 | 1 | 14907 | 14,906 | 06-1 |
| 12 | 06 | 25270246 | 25285963 | 15,717 | 06-3 |
| 13 | 07 | 5 | 21255 | 21,250 | 07-1 |
| 14 | 07 | 30117208 | 30125630 | 8,422 | 07-2 |
| 15 | 08 | 4 | 14837 | 14,833 | 08-1 |
| 16 | 08 | 25034561 | 25046223 | 11,662 | 08-2 |
| 17 | 09 | 4 | 7698 | 7,694 | 09-1 |
| 18 | 09 | 27076851 | 27091241 | 14,390 | 09-2 |
| 19 | 10 | 2 | 22505 | 22,503 | 10-1 |
| 20 | 10 | 29458137 | 29459543 | 1,406 | 10-2 |
| 21 | 11 | 4 | 26556 | 26,552 | 11-1 |
| 22 | 11 | 20604024 | 20615775 | 11,751 | 11-2 |
| 23 | 12 | 3 | 13394 | 13,391 | 12-1 |
| 24 | 12 | 23760205 | 23788023 | 27,818 | 12-2 |
| 25 | 13 | 2 | 16050 | 16,048 | 13-1 |
| 26 | 13 | 29094833 | 29110787 | 15,954 | 13-2 |
| 27 | 14 | 1 | 24525 | 24,524 | 14-1 |
| 28 | 14 | 30173581 | 30183698 | 10,117 | 14-2 |
| 29 | 15 | 6 | 16870 | 16,864 | 15-1 |
| 30 | 15 | 22920453 | 22938019 | 17,566 | 15-2 |
| 31 | 16 | 19 | 6632 | 6,613 | 16-1 |
| 32 | 16 | 26381321 | 26409091 | 27,770 | 16-2 |
| 33 | 17 | 4 | 10257 | 10,253 | 17-1 |
| 34 | 17 | 20994128 | 21002562 | 8,434 | 17-2 |
| 35 | 18 | 1 | 11571 | 11,570 | 18-1 |
| 36 | 18 | 39374073 | 39405660 | 31,587 | 18-2 |
| 37 | 19 | 4 | 25010 | 25,006 | 19-1 |
| 38 | 19 | 27208317 | 27238938 | 30,621 | 19-2 |
| Total |  |  |  | 617,602 |  |
| Mean |  |  |  | 16,253 |  |

These sizes are consistent with the predictions reported for the PN40024 (23). However, our results differ from those described for Thompson Seedless (24). While the authors predicted the longest telomere to be located on chromosome 19 with a length of 23.7 kb, we identified it on one arm of chromosome 1, exceeding 43 kb and comprising 1966 uniformly distributed 21-mer repeats, as shown in the zoomed view of the first sector of this chromosome (**Figure 1B**). For comparison, we observed much shorter telomeres in *A. thaliana* (average length ∼3 kb) (**Figure S2A-C, Table S2**), whereas *Z. mays* displayed considerably longer telomeres (average length ∼74 kb, with three exceeding 100 kb) (**Figure S2D-F, Table S3**).

The visual representation of the telomeric 21-mers along the chromosomes, particularly at the extremities, revealed that the PN40024 genotype telomeric arrays do not always map exactly in correspondence with chromosome ends, but instead are separated by small terminal gaps ranging from 27 to 132 kb. This feature appears to be specific to PN40024 as it was not observed in Thompson Seedless or in the two other plant species analyzed.

Moreover, one telomere seems to be absent from chromosomes 15 and 17 in PN40024, a phenomenon not observed in Sultana. Interestingly, in both cultivars, a small yet consistent block of telomeric repeats was detected in the middle of chromosome 6, suggesting the potential fusion of two ancestral chromosomes.

A similar phenomenon has been observed on chromosome 1 of *A. thaliana* (36, 37), as confirmed by the high number of interstitial telomeric repeats we detected when analyzing the Arabidopsis T2T genome. The relatively high abundance and proportion of telomeric sequences in grapevine cultivars, more than tenfold higher than Arabidopsis and comparable to those in maize, suggest that they represent suitable candidates for proof-of-concept experimental trials. This abundance increased the likelihood of successful multi-binding events, highlighting grapevine as a promising model for the development and optimization of telomere-targeting techniques.

As for tomato, the other model species we chose to explore, we did not consider this verification step to be necessary, since telomeric regions in this species have already been successfully targeted in previous CRISPR-Cas-based applications (21).

### *In vitro* fragment recognition and binding specificity assays of the (d)Cas9 loaded with the telomeric gRNA

After confirming that the grapevine telomeric regions are theoretically long enough to support efficient binding of the RNP complex, we selected the sequence GGGTTTAGGGTTTAGGGTTT as the spacer. This sequence was chosen based on its established use in CRISPR–FISH applications (18, 22). Notably, when this motif occurs in the genome it is immediately followed by an AGG protospacer adjacent motif (PAM), which is required Cas9 recognition of the target sequence. The guide RNA (gRNA) was assembled by combining a CRISPR-encoding RNA (crRNA) containing the spacer sequence with a trans-activating CRISPR RNA (tracrRNA), which together form the structure required Cas protein binding and target recognition. Although the fluorophore on the tracrRNA was not required for telomeric chromatin enrichment, it was initially included to enable imaging-based validation. As a non-targeting control, we also employed a gRNA specific to the *E. coli GAL4* gene (11). The functionality of an RNP assembled with either gRNAs and (d)Cas9 was validated through *in vitro* cleavage and binding assays using amplified DNA fragments containing either a single telomeric or *GAL4* spacer (**Figure 2**).

**Figure 2.**
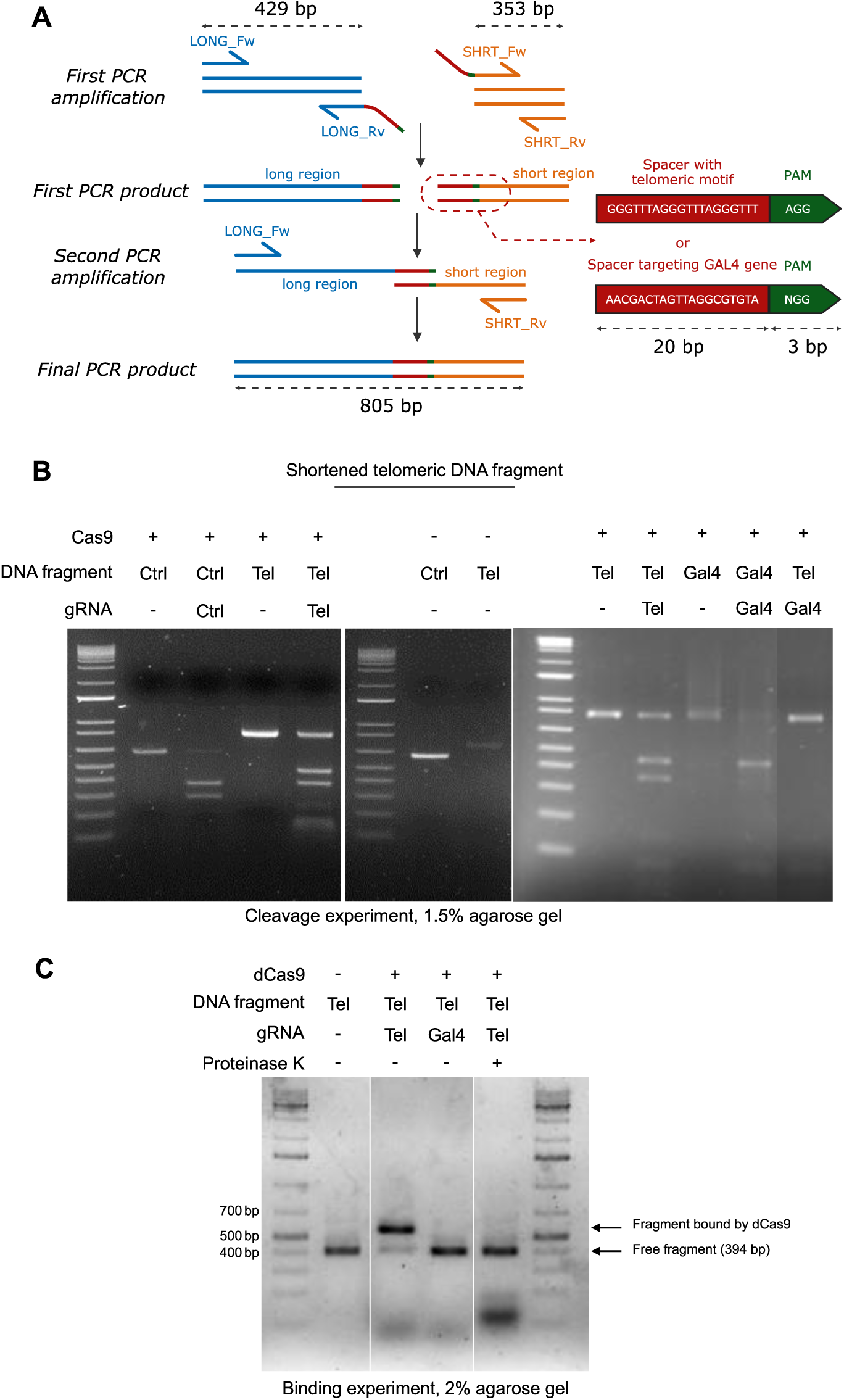
*In vitro* validation of Cas9/dCas9 specificity guided by telomeric gRNA. (**A**) Schematic representation of the PCR amplification strategy used to generate a fragment containing a single telomeric spacer (GGGTTTAGGGTTTAGGGTTT) adjacent to an AGG PAM sequence. Primer positions (LONG_Fw/Rv and SHORT_Fw/Rv) and the resulting amplicons are indicated. (**B**) Cleavage assay. Telomeric (Tel), control (Ctrl) or *GAL4* (Gal4) fragments were incubated with active Cas9 in the presence of the corresponding gRNA, a non-corresponding gRNA, or in the absence of gRNA. Reactions were carried out for 1 h at 37 °C, followed by enzyme inactivation at 80 °C for 5 min, and products were resolved on agarose gel. (**C**) Binding assay. A shortened telomeric fragment (394 bp), obtained using internal primers, was incubated with catalytically inactive dCas9 and either the telomeric gRNA or a *GAL4* gRNA, in the presence or absence of proteinase K. Complex formation was assessed by electrophoretic mobility shift assay (EMSA) on a 2% agarose gel.

### CRISPR–FISH on telomeres allows the observation of foci on nuclei from grapevine and tomato

To verify that the dCas9–gRNA ribonucleoprotein (RNP) complex could access and bind its target sequences within fixed nuclei, we applied a telomere repeat-specific CRISPR–FISH protocol to nuclei of our model species. This step was necessary both to assess the accessibility of the RNP complex to chromatin in fixed nuclei and to confirm that dCas9 could efficiently bind telomeric regions, which are known to display an intermediate chromatin state between euchromatin and heterochromatin. The CRISPR–FISH protocol, whose efficiency and specificity have previously been demonstrated (18), was applied to *V. vinifera* cv. Thompson Seedless and to tomato of Micro-Tom varieties. In both cases, we detected clear and distinct putative telomeric *foci* (**Figure 3A,E**). To confirm the telomeric origin of these signals, we combined CRISPR–FISH with conventional DNA–FISH, enabling direct comparison of their nuclear localization. Because the CRISPR–FISH signals weakened after DNA-FISH, we first recorded the coordinates of nuclei exhibiting CRISPR–FISH signals, then proceeded with DNA-FISH and imaged the same nuclei (**Figure 3B,F**). The images obtained by the two approaches were then manually aligned using the corresponding DAPI channels (**Figure 3C,G**) revealing a clear colocalization of telomere foci detected by both CRISPR-FISH and DNA-FISH methods (**Figure 3D,H**). Indeed, most CRISPR–FISH signals corresponded to a FISH signals. Additional CRISPR-FISH and DNA-FISH assays performed independently in Thompson Seedless, Cabernet franc, and tomato confirmed these observations (**Figure S3**). In all cases, the number of foci detected per nucleus was lower than the expected number of telomeres, a discrepancy likely due to factors such as nuclear three-dimensional architecture, chromatin accessibility, or dCas9 binding efficiency. Notably, CRISPR–FISH consistently revealed a greater number of foci compared to DNA–FISH. As expected, no nuclear foci were observed when CRISPR-FISH was performed using a gRNA targeting the bacterial *GAL4* gene or when the dCas9 was loaded only with the ATTO-labeled tracrRNA (**Figure S4**).

**Figure 3.**
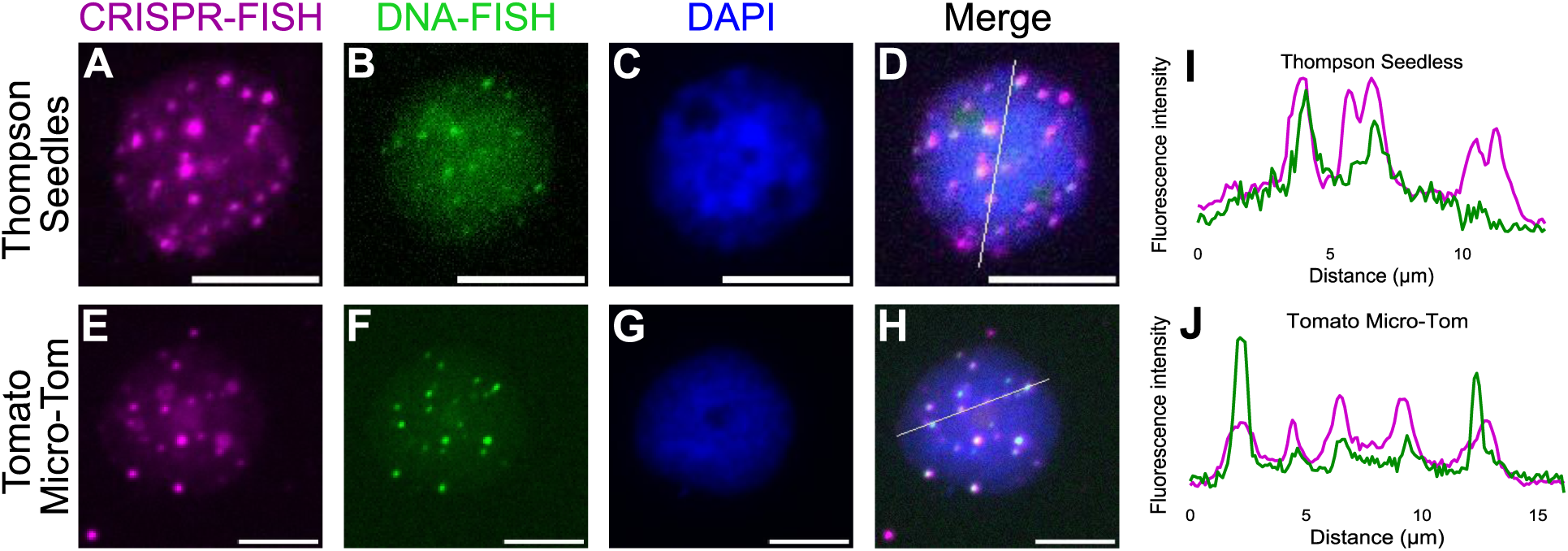
Colocalization of telomeric signals detected by CRISPR-FISH and DNA-FISH in plant nuclei. Representative nuclei from *V. vinifera* cv. Thompson Seedless (**A–D**) and tomato (*S. lycopersicum* cv. Micro-Tom; **E–H**) are shown. CRISPR-FISH signals (magenta; **A, E**), DNA-FISH signals (green; **B, F**), and DAPI staining (blue; **C, G**) are presented separately and as merged images (**D, H**). Line-scan fluorescence intensity profiles along the indicated axes in merged images are shown for Thompson Seedless (**I**) and Micro-Tom (**J**), highlighting the spatial overlap between CRISPR-FISH and DNA-FISH signals. Scale bars: 8 µm.

Here, CRISPR-FISH and FISH provided a clear, visual and straightforward way to evaluate the target effectiveness in nuclei of grapevine and tomato. This technique could very well benefit from further refinement such as quantitative images analyses and optimization of the protocols for simultaneous labeling of FISH and CRISPR-FISH, the application of CRISPR-FISH served primarily as a validation step to assess the suitability of CRISPR-dCas9 system in purified grapevine and tomato nuclei, therefore further improvements of the technique are beyond the scope of this paper. Furthermore, while the imaging method is relatively quick to set up and execute, its applicability may be limited to specific species and target sequences, as it strongly depends on the abundance of sequence repetition.

Overall, the overlap between CRISPR–FISH and DNA–FISH signals support the preservation of a chromatin organization compatible with dCas9 targeting in isolated nuclei, while avoiding the major perturbations associated with protoplast-based transformation approaches. Although the chromatin architecture observed in isolated nuclei may differ to some extent from that of intact tissues, this system represents a robust and physiologically relevant compromise for investigating DNA–protein interactions under controlled conditions.

### Assessment of the transfection rate of purified grapevine and tomato nuclei

After validating the ability of the RNP complex to target telomeric sequences in formaldehyde-fixed nuclei spread on microscope slides, we next asked whether the RNP complex could also access isolated nuclei directly in solution. This step was essential to determine whether purified nuclei could efficiently internalize the RNP complex, a prerequisite for performing locus-specific chromatin precipitation without relying on cellular transformation. To address this, we adapted a nuclear extraction protocol and evaluated the capacity of purified nuclei to internalize the RNP complex. Fluorescence analyses performed using a CellDrop cell counter to assess transfection efficiency revealed mean uptake rates of 68% in grapevine nuclei and 38% in tomato nuclei, with fluorescence distributed throughout the entire nucleus. Comparable, although slightly lower, values were obtained when using either a telomere-targeting gRNA or a control gRNA specific for the *Escherichia coli GAL4* gene (11), with uptake rates of 65% for grapevine and 28% for tomato. These results indicate that the assay primarily measures the overall uptake of the fluorescently labeled RNP complex into the nucleus rather than its specific binding to the target sequence. Varying the number of nuclei in the transfection (from 10^7^ to 10^8^) did not significantly affect efficiency in grapevine, whereas it affected efficiency in tomato. Therefore, for grapevine, we considered the counts from samples obtained by transfecting both 10^7^ and 10^8^ nuclei, whereas for tomato, we retained only the 10^8^ nuclei samples (**Figure S5**; **Table S4**).

### Precipitation of telomeric regions and sequence analysis

After confirming by CRISPR–FISH that dCas9 can specifically bind telomeric sequences in paraformaldehyde-fixed nuclei, we next assess whether this approach could be utilized to precipitate telomeric chromatin. For this purpose, we adapted a protocol previously developed in mammalian cells (11). Nuclei suspensions from grapevine and tomato leaves were incubated for 1-2 hours with a biotinylated dCas9 loaded with the telomere-specific Alt-R guide. Following incubation, nuclei were fixed with formaldehyde, and chromatin was extracted and fragmented by sonication. The biotinylated dCas9 was then precipitated using streptavidin-coated magnetic beads, and the recovered DNA fragments were sequenced with Illumina technology.

Because of the repetitive nature of telomeric sequences standard quantification methods were not applicable: 21-mer, qPCR could not be applied due to the difficulty in primer or probes design, and sequence reads could not be reliably mapped to the reference genome. Therefore, we assessed the content of the precipitated DNA by identifying and counting the 21-mers present in the sequencing output. A control experiment was performed using dCas9 loaded with *GAL4*-specific gRNA. An increase in the frequency of telomeric 21-mers relative to other 21-mers was observed in samples precipitated with the telomere-specific gRNA compared with both the non-precipitated input DNA and the *GAL4* control, while non-telomeric 21-mers remained at background levels. This enrichment was consistently observed in all tested genotypes, including *V. vinifera* cv. Thompson Seedless, *V. vinifera* cv. Cabernet Franc, and *S. lycopersicum* cv. Micro-Tom, although with differences in magnitude among species and cultivars. Notably, telomeric 21-mers showed enrichment values several orders of magnitude higher than non-telomeric sequences, demonstrating both the specificity and efficiency of the dCas9-mediated chromatin precipitation approach. (**Figure 4**).

**Figure 4.**
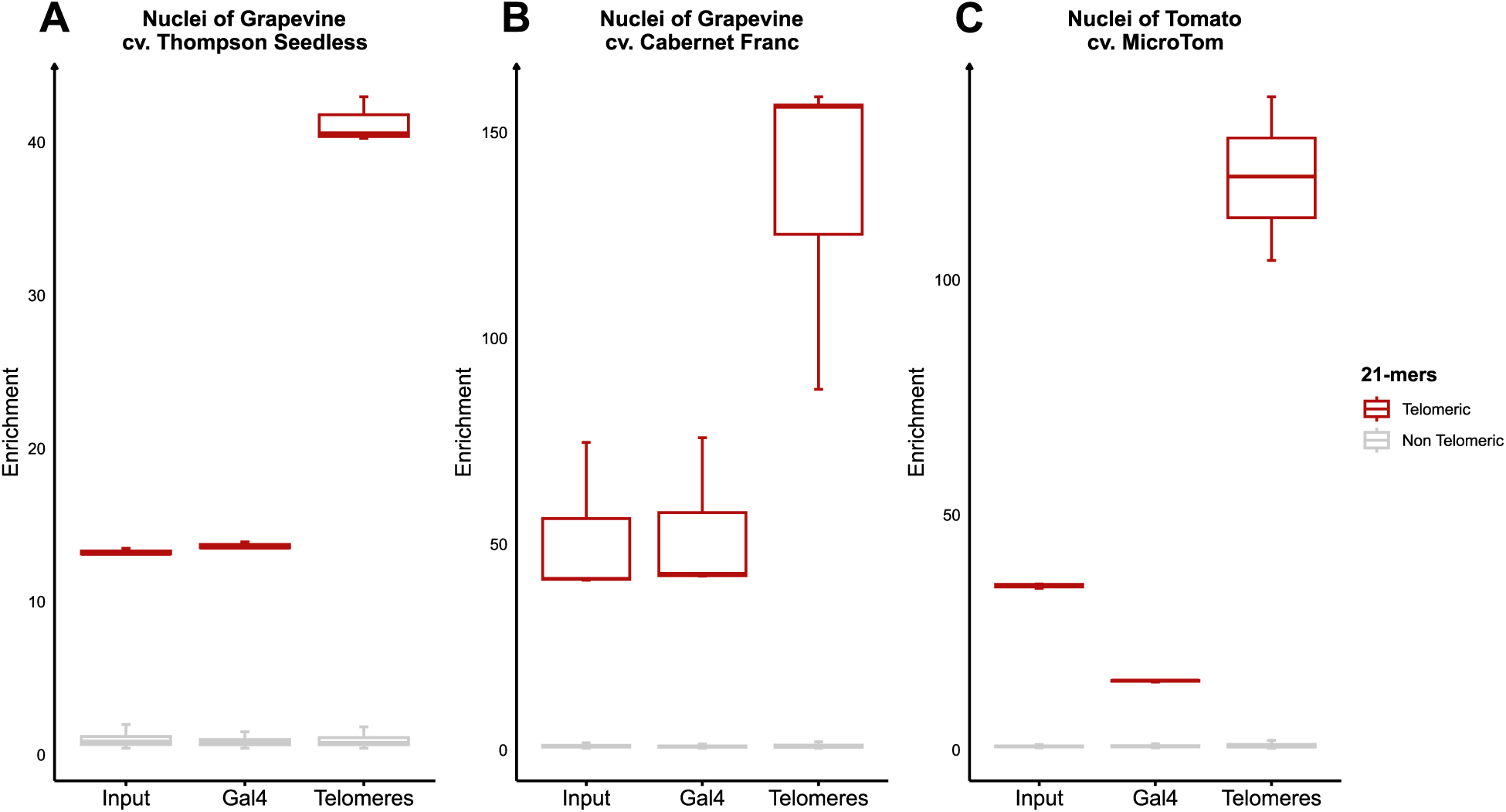
Telomere enrichment following CRISPR-based chromatin precipitation in grapevine and tomato. Leaf nuclei from *V. vinifera* cv. Thompson Seedless (**A**), cv. Cabernet Franc **(B),** and *S. lycopersicum* cv. MicroTom **(C)** were isolated and incubated with a biotinylated dCas9 loaded either with telomere-specific gRNA or with *GAL4* gRNA. Chromatin was subsequently captured using streptavidin-coated beads, and the recovered DNA (precipitated fraction) was sequenced on an Illumina platform. In parallel, a non-precipitated fraction of each sample was sequenced as control. The abundance of telomeric and non-telomeric 21-mers was quantified, and enrichment values were calculated as the ratio of each 21-mer count to the mean count of all 21-mers within the same sample. Box plots show the distribution of enrichment values for telomeric (red) and non-telomeric (grey) 21-mers.

### Precipitation of low copy and single-copy loci

After demonstrating the feasibility of the protocol on highly repeated loci through NGS sequencing of the precipitated fraction, we next aimed to target more typical genomic regions whose spacers occurs only a few times, or even once, in the genome. To avoid directly targeting strictly single-copy loci, which could increase the risk of obtaining insufficient precipitated material, we first searched for sequences whose spacers targets were present in multiple copies. This led us to focus on the stilbene synthase gene family, which is specific to certain plants species, including grapevine, and displays a high degree of sequence homology among its members (38). We therefore selected two spacers located in the promoter regions of different members of the family: one present 18 times in the genome (TAGAAACGCTCAACGTGCCA, hereafter referred to as gRNA STS #1) (**Figure 5A,C**) and another present only twice (TTACATGGGCTAAAGACAAA, gRNA STS #2) (**Figure 5A,B**). In the same experiment, we also designed a spacer targeting the unique grapevine *VvACTIN* gene (**Figure 5A**, gRNA for *VvACTIN*). In this case, the different gRNAs were not assembled from two separate components as for telomeric targets, but were *in vitro* transcribed using a T7 polymerase, as single guide RNAs (sgRNA) from a DNA template containing the T7 promoter and encoding the desired spacer together with a scaffold previously described in literature as efficient (12). Once obtained, the sgRNAs were tested in cleavage assays as described before. For this purpose, the cleaved fragments were amplified as describe for the telomeric spacer and the *GAL4* spacer for gRNA #1 and #2, whereas amplification for the *VvACTIN* gRNA was performed directly from genomic DNA, since this target is not present in repeated copies. All gRNAs produced positive results in the cleavage assays (**Figure 5D-E**); therefore, the precipitation experiments were subsequently performed.

**Figure 5.**
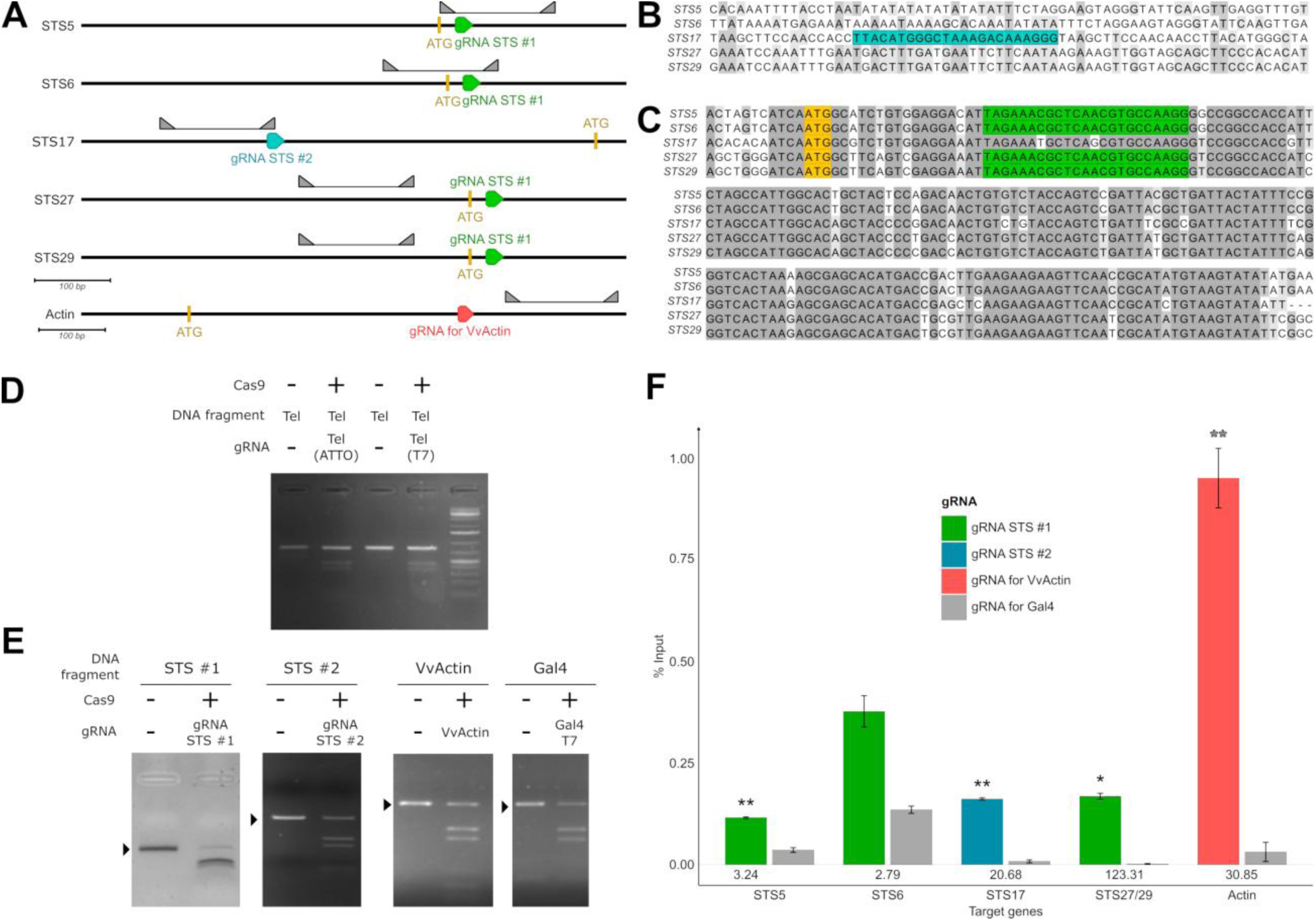
Targeted precipitation of low-copy genomic regions using CRISPR-based chromatin capture. (**A**) Genomic positions of gRNA target sites within selected *STS* family genes and *VvACTIN* in the *V. vinifera* cv. Thompson Seedless genome, together with the positions of qPCR primers. Gene sequences were retrieved from the reference genome and visualized in Geneious software. Scale bars are shown below each panel. (**B-C**) Sequence alignment of selected *STS* genes (*STS5*, *STS6*, *STS17*, *STS27*, and *STS29*) at the regions targeted by the designed spacers. (**B**) Sequence alignment of promoter regions including the gRNA STS #1 target site. The region targeted by gRNA STS #2 is highlighted in blue. (**C**) Alignment of the region spanning from −12 bp to +191 bp relative to the ATG start codon; the region targeted by gRNA STS #2 is highlighted in green. Grey shading indicates sequence identity across aligned sequences. (**D**) *In vitro* cleavage assay performed on PCR-amplified fragments containing gRNA target sites for telomeres, using Cas9 in the presence or absence of gRNA. (**F**) *In vitro* cleavage assays on PCR-amplified fragments corresponding to STS #1, STS #2, *VvACTIN*, and *GAL4* target regions, using Cas9 in the presence of the corresponding gRNAs. Arrowheads indicate cleavage products. (**E**) Quantification of chromatin precipitation efficiency. Nuclei isolated from V. vinifera cv. Thompson Seedless were incubated with biotinylated dCas9 loaded with gRNA STS #1, gRNA STS #2, gRNA targeting VvACTIN, or GAL4 gRNA as negative control. qPCR was performed on target loci (STS5, STS6, STS17, STS27/29, and Actin) in both precipitated and input fractions. Enrichment is expressed as percentage of input (% input). Colored bars represent specific gRNA targeting, whereas grey bars correspond to GAL4 controls. Values below the bars indicate fold enrichment relative to GAL4. Statistical significance was assessed using Student’s t-test (P < 0.05; P < 0.01).

The precipitations were carried out on nuclei isolated from the grapevine cultivar Thompson Seedless, following the procedure described above. Each transfection was performed starting from 10^8^ nuclei per transfection and corresponded to a ribonucleoprotein composted of biotinylated dCas9 and a single gRNA species. After precipitation or recovery of input fraction, DNA from both precipitated and input samples was recovered in a final volume of 50 µL and subsequently analyzed by qPCR. The primers used for qPCR amplification were designed in Geneious software on the sequence of the target gene (or its promoter, when appropriate) so as to amplify a ∼150 pb genomic fragment located within a ±300 pb around the spacer site. qPCR was chosen as analytical method because it allowed rapid evaluation of target enrichment immediately after the precipitation step. For each gRNA, the enrichment of several selected targets loci was evaluated by comparing precipitated samples with their corresponding inputs. For gRNA #1, enrichment was evaluated for the promoters of *STS5*, *STS6* and *STS27/29* (indistinguishable at the amplified position), while *STS17* was analyzed for gRNA #2 (**Figure 5F**).

The precipitation of the same genes was also evaluated in samples precipitated with the *GAL4* gRNA control. After analysis of the CT values, and normalization on the corresponding input samples, enrichment of the target sequences was expressed as percentage of input. For most target genes (*STS5*, *STS17*, *STS27/29* and *VvACTIN*), enrichment obtained with the corresponding gRNA was consistently significatively higher to that observed with the *GAL4* gRNA control. This was observed both for repeated loci, which may benefit from the simultaneous precipitation of multiple genomic regions, and for the single-copy *VvACTIN* locus. For *STS6*, a higher enrichment was observed with the specific gRNA compared to the *GAL4* control; however, this difference was not statistically significant (p > 0.05). The lack of statistical significance is likely due to high variability among biological replicates.

The enrichment obtained with the specific gRNA was also compared with that obtained using the telomeric gRNA, by performing qPCR on the same target genes (**Figure S6**). As expected, enrichment values were generally higher in samples precipitated with the corresponding specific gRNAs. However, in some cases, significative differences were also observed between precipitations performed with the *GAL4* gRNA and telomeric gRNA. One possible explanation is that the telomeric RNP complex, which binds to highly repetitive telomeric sequences, may remain associated with chromatin for longer periods compared with the *GAL4* RNP, which lacks a genomic target in grapevine. This prolonged chromatin residence could increase the likelihood of co-precipitating genomic regions that are spatially proximal to telomeres within the three-dimensional organization of the nucleus. In contrast, the *GAL4*-guided RNP is unlikely to stably associate with chromatin and may therefore display lower levels of background precipitation. Interestingly, the level of non-specific precipitation observed with the *GAL4* and telomeric gRNAs appeared relatively constant across samples, whereas target-specific enrichments displayed greater variability. This variability likely reflects differences in gRNA efficiency and target accessibility, which may depend on factors such as chromatin context, PAM sequence composition, and spacer–target pairing efficiency (39, 40). A notable difference should also be considered between the single-copy *VvACTIN* spacer and *STS* spacers targeting repeated loci. In addition to potential differences in gRNA specificity, the number of genomic targets may also influence the observed enrichment. While the RNP complex loaded with the *VvACTIN* gRNA is directed to a single genomic locus, an RNP guided by spacers targeting repeated loci may bind to multiple genomic targets, thereby distributing the precipitated material across different loci and reducing the apparent enrichment measured for each individual gene. Consequently, the measured enrichment likely reflects the combined effects of gRNA efficiency, target copy number, and locus accessibility, together with a minor contribution of technical variability between samples, which may account for the discrepancies observed among the precipitated target genes. In contrast with the initial imaging trials, chromatin precipitation followed by sequencing provides a more direct and universal approach for assessing specificity, as it relies on the actual nucleotide sequence identity of the precipitated fragments rather than on sequence abundance. Moreover, the precipitation step, performed prior to reverse-crosslinking, provides access to a purified chromatin fraction of the genome, thereby enabling additional downstream analyses with only minor modifications to the protocol. In the studies that inspired our work, this approach was successfully used to identify proteins associated to the targeted loci through proteomic analysis (11, 12) and to investigate long-range chromatin interactions (11). Establishing a comparable protocol in plants therefore represents a natural continuation and expansion of the present work.

Taken together, these results demonstrate the feasibility of precipitating the DNA fraction of specific chromatinic regions using biotinylated CRISPR-dCas9, even for targets present as single copy loci in the genome. They also highlight the high target specificity of dCas9 for its target sequences. One of the main challenges for implementing the downstream applications mentioned above is the amount of starting material required. In this study, we showed that transfection of ∼10^7^ grapevine nuclei, with an uptake efficiency of ∼70%, was sufficient to recover DNA in quantities compatible with sequencing analyses. However, proteomic applications will likely demand higher yields, which may necessitate further optimization of either transfection parameters or chromatin extraction protocols. For studies of gene regulation, a key advantage of our system is that the dCas9 RNP complex is introduced into purified, fixed nuclei rather than into living cells. This approach minimizes potential artifacts such as changes in gene expression induced by dCas9 binding or the detection of non-nuclear interactors (12). Indeed, in this system, the experimental target is no longer the living organism but its purified nuclei, fixed at a defined physiological state prior to the introduction of any exogenous components and, if desired, after the application of a specific stress to investigate the regulation of responsive genes. Under these conditions, it is reasonable to assume that gene expression is influenced solely by the applied stress. Transfection nevertheless remains a critical step and relies on protocols that are often species- and tissue-specific. Although several transformation techniques are available, none of them are universal and many plant species remain recalcitrant to transformation (41). For this reason, we aimed to develop a protocol that circumvents the transformation step, providing a major advantage in terms of transferability across species, genotypes, tissues, and experimental contexts, with only modest adjustments to nuclei isolation procedures, which are far less demanding than the development or optimization of transformation systems.

## CONCLUSIONS

Our results demonstrate the feasibility of specifically targeting both repetitive and non-repetitive DNA sequences in purified plant nuclei by exploiting the precise recognition capabilities of the CRISPR-dCas9 ribonucleoprotein complex. We validated the affinity of dCas9 for its target sequence through *in vitro* cleavage assays and confirmed its specificity in plant nuclei using both imaging-based and genomic approaches.

There is an increasing demand for techniques that enable the study of chromatin at defined genomic loci across diverse eukaryote systems. Here, we describe GRASP, a protocol for chromatin extraction from isolated nuclei of two plant species, grapevine and tomato, and the selective capture of their telomeric chromatin fraction, with specificity confirmed by sequence-enrichment analysis on the precipitated DNA. The use of a biotinylated dCas9-gRNA complex enables antibody-independent chromatin capture, thereby avoiding limitations related to antibody availability, specificity and efficiency. Our choice to work with isolated nuclei rather than other systems, such as protoplasts, provides several advantages. Nuclear isolation is simpler, faster, and yields a higher number of “transfectable units”. In addition, nuclei extracted from formaldehyde-fixed tissues preserve the native chromatin organization of the original cells, whereas protoplast isolation requires enzymatic digestion and mechanical treatments, which can induce substantial chromatin remodeling and introduce potential artifacts. Furthermore, nuclei can be isolated from a wide range of plant tissues with minimal protocol adjustments, making this approach broadly adaptable across species and experimental contexts.

By enabling the specific capture of telomeric and single-loci chromatin, this work provides a nucleus-based framework for target chromatin investigations in plants. This strategy opens the way to future analyses of protein‒DNA interactions, chromatin architecture, and genome organization at specific regions. The ability to unravel protein‒DNA interactions at defined genomic locations will be instrumental for understanding the molecular mechanisms of gene regulation, the establishment of specialized chromatin states, and long-range chromatin interactions, as already demonstrated in mammalian systems. Extending such approaches to plants will expand opportunities for genome-wide studies and for investigating chromatin dynamics under diverse developmental, physiological and environmental conditions.

## ACKNOWLEDGEMENT

We are thankful to Pr. Wang who kindly provided us with the AtU6 vector used as template and model for the gRNA and fragment amplification.

## DATA AVAILABILITY

The data that support the findings of this study are openly available in the SRA (Sequence Read Archive) database at https://www.ncbi.nlm.nih.gov/sra under the BioProject ID PRJNA1223867.

## AUTHORS’ CONTRIBUTIONS

Aurélien Devillars: Formal analysis, Methodology, Validation, Writing - original draft. Silvia Farinati: Conceptualization, Formal analysis, Writing - review & editing. Adriana F. Soria Garcia & Justin Joseph: Formal analysis, Writing - review & editing. Sara Zenoni, Edoardo Bertini, & Alessandra Amato: Provision of *in vitro* plant material, Writing - review & editing. Giovanni Gabelli: Formal analysis, Writing - review & editing. Bhanu P. Potlapalli & Andreas Houben: Formal analysis, Writing - review & editing. Fabio Palumbo & Gianni Barcaccia: Writing - review & editing. Alessandro Vannozzi: Conceptualization, Methodology, Validation, Writing - original draft.

## SUPPLEMENTARY MATERIALS

**Figure S1** Abundance of telomeres in the T2T genomes of *V. vinifera* cv Pinot Noir.

**Figure S2** Abundance of telomeres in the T2T genomes of two non-*Vitis* species.

**Figure S3** CRISPR-FISH and Standard FISH on grapevine and tomato nuclei

**Figure S4** CRISPR–FISH on grapevine and tomato nuclei including non-targeting controls.

**Figure S5** Nuclei isolation from grapevine leaves, transformation and counting.

**Figure S6** Precipitation of lesser-repeated spacers with telomeric gRNA as secondary control.

**Table S1** Telomeres of *V. vinifera* cv. Pinot Noir T2T genome assembly

**Table S2** Telomeres *of A. thaliana* T2T genome assembly

**Table S3** Telomeres of *Z. mays* T2T genome assembly

**Table S4** Isolated nuclei transfection rates

**Supplementary Method 1** Reagents and Solutions

**Supplementary Method 2** Kits and specific materials

**Supplementary Method 4** Solutions

**Supplementary Method 4** Methods

**Supplementary Method 5** Troubleshooting points

**Supplementary File 1** Bashscript:Jellyfish_blast_Rgraphs_script and Rscript: chr_k21_telomeric_GENERAL

**Supplementary File 2** Primers and fragment sequences

## REFERENCES

1. Vergara, Z. and Gutierrez, C. (2017) Emerging roles of chromatin in the maintenance of genome organization and function in plants. Genome Biol, 18, 96.

2. Orphanides, G. and Reinberg, D. (2002) A Unified Theory of Gene Expression. Cell, 108, 439–451.

3. Morrison, O. and Thakur, J. (2021) Molecular Complexes at Euchromatin, Heterochromatin and Centromeric Chromatin. Int J Mol Sci, 22, 6922.

4. Achrem, M., Szućko, I. and Kalinka, A. (2020) The epigenetic regulation of centromeres and telomeres in plants and animals. Comp Cytogenet, 14, 265–311.

5. Smith, E.M., Pendlebury, D.F. and Nandakumar, J. (2020) Structural biology of telomeres and telomerase. Cell Mol Life Sci, 77, 61–79.

6. Xie, Y., Liu, Y.-G. and Chen, L. (2016) Assessing protein-DNA interactions: Pros and cons of classic and emerging techniques. Sci. China Life Sci., 59, 425–427.

7. Visa, N. and Jordán-Pla, A. eds (2018) Chromatin Immunoprecipitation: Methods and Protocols Springer New York, New York, NY.

8. Pickar-Oliver, A. and Gersbach, C.A. (2019) The next generation of CRISPR-Cas technologies and applications. Nat Rev Mol Cell Biol, 20, 490–507.

9. Devillars, A., Magon, G., Pirrello, C., Palumbo, F., Farinati, S., Barcaccia, G., Lucchin, M. and Vannozzi, A. (2024) Not Only Editing: A Cas-Cade of CRISPR/Cas-Based Tools for Functional Genomics in Plants and Animals. International Journal of Molecular Sciences, 25, 3271.

10. Schmidtmann, E., Anton, T., Rombaut, P., Herzog, F. and Leonhardt, H. (2016) Determination of local chromatin composition by CasID. Nucleus, 7, 476–484.

11. Liu, X., Zhang, Y., Chen, Y., Li, M., Shao, Z., Zhang, M.Q. and Xu, J. (2018) CAPTURE: *In Situ* Analysis of Chromatin Composition of Endogenous Genomic Loci by Biotinylated dCas9. Current Protocols in Molecular Biology, 123, e64.

12. Wang, Z., He, Z., Liu, Z., Qu, M., Gao, C., Wang, C. and Wang, Y. (2023) A reverse chromatin immunoprecipitation technique based on the CRISPR–dCas9 system. Plant Physiology, 191, 1505–1519.

13. Qi, L.S., Larson, M.H., Gilbert, L.A., Doudna, J.A., Weissman, J.S., Arkin, A.P. and Lim, W.A. (2013) Repurposing CRISPR as an RNA-Guided Platform for Sequence-Specific Control of Gene Expression. Cell, 152, 1173–1183.

14. Veena, V., Jiang, H., Doerge, R. w. and Gelvin, S.B. (2003) Transfer of T-DNA and Vir proteins to plant cells by Agrobacterium tumefaciens induces expression of host genes involved in mediating transformation and suppresses host defense gene expression. The Plant Journal, 35, 219–236.

15. Hu, Y., Lacroix, B. and Citovsky, V. (2021) Modulation of plant DNA damage response gene expression during Agrobacterium infection. Biochemical and Biophysical Research Communications, 554, 7–12.

16. Procházková Schrumpfová, P., Fojtová, M. and Fajkus, J. (2019) Telomeres in Plants and Humans: Not So Different, Not So Similar. Cells, 8, 58.

17. Dreissig, S., Schiml, S., Schindele, P., Weiss, O., Rutten, T., Schubert, V., Gladilin, E., Mette, M.F., Puchta, H. and Houben, A. (2017) Live-cell CRISPR imaging in plants reveals dynamic telomere movements. The Plant Journal, 91, 565–573.

18. Ishii, T., Schubert, V., Khosravi, S., Dreissig, S., Metje-Sprink, J., Sprink, T., Fuchs, J., Meister, A. and Houben, A. (2019) RNA-guided endonuclease – in situ labelling (RGEN-ISL): a fast CRISPR/Cas9-based method to label genomic sequences in various species. New Phytologist, 222, 1652–1661.

19. Němečková, A., Wäsch, C., Schubert, V., Ishii, T., Hřibová, E. and Houben, A. (2019) CRISPR/Cas9-Based RGEN-ISL Allows the Simultaneous and Specific Visualization of Proteins, DNA Repeats, and Sites of DNA Replication. CGR, 159, 48–53.

20. Potlapalli, B.P., Schubert, V., Metje-Sprink, J., Liehr, T. and Houben, A. (2020) Application of Tris-HCl Allows the Specific Labeling of Regularly Prepared Chromosomes by CRISPR-FISH. Cytogenet Genome Res, 160, 156–165.

21. Nagaki, K. and Yamaji, N. (2020) Decrosslinking enables visualization of RNA-guided endonuclease–in situ labeling signals for DNA sequences in plant tissues. Journal of Experimental Botany, 71, 1792–1800.

22. Khosravi, S., Schindele, P., Gladilin, E., Dunemann, F., Rutten, T., Puchta, H. and Houben, A. (2020) Application of Aptamers Improves CRISPR-Based Live Imaging of Plant Telomeres. Front Plant Sci, 11, 1254.

23. Shi, X., Cao, S., Wang, X., Huang, S., Wang, Y., Liu, Z., Liu, W., Leng, X., Peng, Y., Wang, N., et al. (2023) The complete reference genome for grapevine (*Vitis vinifera* L.) genetics and breeding. Horticulture Research, 10, uhad061.

24. Wang, X., Tu, M., Wang, Y., Zhang, Y., Yin, W., Fang, J., Gao, M., Li, Z., Zhan, W., Fang, Y., et al. (2024) Telomere-to-telomere and gap-free genome assembly of a susceptible grapevine species (Thompson Seedless) to facilitate grape functional genomics. Horticulture Research, 11, uhad260.

25. Fulcher, N., Teubenbacher, A., Kerdaffrec, E., Farlow, A., Nordborg, M. and Riha, K. (2015) Genetic Architecture of Natural Variation of Telomere Length in Arabidopsis thaliana. Genetics, 199, 625–635.

26. Chen, J., Wang, Z., Tan, K., Huang, W., Shi, J., Li, T., Hu, J., Wang, K., Wang, C., Xin, B., et al. (2023) A complete telomere-to-telomere assembly of the maize genome. Nat Genet, 10.1038/s41588-023-01419-6.

27. Potlapalli, B.P., Fuchs, J., Rutten, T., Meister, A. and Houben, A. (2024) The potential of ALFA-tag and tyramide-based fluorescence signal amplification to expand the CRISPR-based DNA imaging toolkit. Journal of Experimental Botany, 10.1093/jxb/erae341.

28. Kuo, Y.-T., Câmara, A.S., Schubert, V., Neumann, P., Macas, J., Melzer, M., Chen, J., Fuchs, J., Abel, S., Klocke, E., et al. (2023) Holocentromeres can consist of merely a few megabase-sized satellite arrays. Nat Commun, 14, 3502.

29. Choi, J.Y., Abdulkina, L.R., Yin, J., Chastukhina, I.B., Lovell, J.T., Agabekian, I.A., Young, P.G., Razzaque, S., Shippen, D.E., Juenger, T.E., et al. (2021) Natural variation in plant telomere length is associated with flowering time. The Plant Cell, 33, 1118–1134.

30. Burr, B., Burr, F.A., Matz, E.C. and Romero-Severson, J. (1992) Pinning down loose ends: mapping telomeres and factors affecting their length. The Plant Cell, 4, 953–960.

31. Ev, S. and De, S. (2004) Length regulation and dynamics of individual telomere tracts in wild-type Arabidopsis. The Plant cell, 16.

32. Shakirov, E.V., Salzberg, S.L., Alam, M. and Shippen, D.E. (2008) Analysis of Carica papaya Telomeres and Telomere-Associated Proteins: Insights into the Evolution of Telomere Maintenance in Brassicales. Tropical Plant Biol., 1, 202–215.

33. Kilian, A., Stiff, C. and Kleinhofs, A. (1995) Barley telomeres shorten during differentiation but grow in callus culture. Proceedings of the National Academy of Sciences, 92, 9555–9559.

34. Andreu-Sánchez, S., Aubert, G., Ripoll-Cladellas, A., Henkelman, S., Zhernakova, D.V., Sinha, T., Kurilshikov, A., Cenit, M.C., Jan Bonder, M., Franke, L., et al. (2022) Genetic, parental and lifestyle factors influence telomere length. Commun Biol, 5, 565.

35. Rabanal, F.A., Gräff, M., Lanz, C., Fritschi, K., Llaca, V., Lang, M., Carbonell-Bejerano, P., Henderson, I. and Weigel, D. (2022) Pushing the limits of HiFi assemblies reveals centromere diversity between two Arabidopsis thaliana genomes. Nucleic Acids Res, 50, 12309–12327.

36. Koch, M.A. and Kiefer, M. (2005) Genome evolution among cruciferous plants: a lecture from the comparison of the genetic maps of three diploid species— *Capsella rubella, Arabidopsis lyrata subsp. petraea*, and *A. thaliana*. American J of Botany, 92, 761–767.

37. Kawabe, A., Hansson, B., Hagenblad, J., Forrest, A. and Charlesworth, D. (2006) Centromere Locations and Associated Chromosome Rearrangements in Arabidopsis lyrata and A. thaliana. Genetics, 173, 1613–1619.

38. Vannozzi, A., Dry, I.B., Fasoli, M., Zenoni, S. and Lucchin, M. (2012) Genome-wide analysis of the grapevine stilbene synthase multigenic family: genomic organization and expression profiles upon biotic and abiotic stresses. BMC Plant Biol, 12, 130.

39. Doench, J.G., Fusi, N., Sullender, M., Hegde, M., Vaimberg, E.W., Donovan, K.F., Smith, I., Tothova, Z., Wilen, C., Orchard, R., et al. (2016) Optimized sgRNA design to maximize activity and minimize off-target effects of CRISPR-Cas9. Nat Biotechnol, 34, 184–191.

40. Liang, G., Zhang, H., Lou, D. and Yu, D. (2016) Selection of highly efficient sgRNAs for CRISPR/Cas9-based plant genome editing. Sci Rep, 6, 21451.

41. Levengood, H., Zhou, Y. and Zhang, C. (2024) Advancements in plant transformation: from traditional methods to cutting-edge techniques and emerging model species. Plant Cell Rep, 43, 273.

