## Supplementary material for "GRASP: A PLANT TRANSFORMATION-INDEPENDENT CRISPR-BASED SYSTEM FOR AFFINITY PURIFICATION OF SPECIFIC CHROMATIN LOCI": Supplemetary Methods

Supplementary method 1: Reagents

- AltR ® CRISPR Cas9 crRNA with telomeric spacer for plant telomeres, Sequence: "GGGUUUAGGGUUUAGGGUUUGUUUUAGAGCUAUGCU" (Integrated DNA Technologies, cat. no. NA)
- Alt-R® CRISPR-Cas9 tracrRNA, ATTO550 (Integrated DNA Technologies, cat. no. 1075928)
- Calcium chloride dihydrate (CaCl2.2H2O) (Duchefa Biochemie, cat. no. C0504)
- Cefotaxime sodium (Duchefa Biochemie, cat. no. C0111)
- Chloroform (Sigma-Aldrich®, cat. no. 650498)
- cOmplete™, Mini, EDTA-free Protease Inhibitor Cocktail (Roche™, cat. no. 4693159001)
- dCas9-3XFLAG-Biotin (dCas9) (Sigma-Aldrich®, cat. no. DCAS9PROT)
- Deionized formamide (Millipore®, cat. no. 4610-OP)
- Dextrane (Thermo Scientific Chemicals™, cat. no. 406275000)
- Di-Sodium hydrogen phosphate dihydrate (Na2HPO4.2H2O) (Duchefa Biochemie, cat. no. S0537)
- Dithiothreitol (DTT) (Thermo Scientific™, cat. no. R0861)
- DNA-FISH Probe (, cat. no. )
- DNase-free RNase A (Thermo Scientific™, cat. no. EN0531)
- EDTA (Sigma-Aldrich®, cat. no. E6758)
- EDTA disodium dihydrate (Na2-EDTA) (Duchefa Biochemie, cat. no. E0511)
- Formaldehyde (Sigma-Aldrich®, cat. no. F8775)
- Glycerol (Sigma-Aldrich®, cat. no. G5516)
- Glycine (Sigma-Aldrich®, cat. no. G5417)
- HEPES (Sigma-Aldrich®, cat. no. PHG0001)
- Indole-3-butyric acid (IBA) (Duchefa Biochemie, cat. no. I0902)
- Isoamyl alcohol (Sigma-Aldrich®, cat. no. W205702)
- Lithium chloride (LiCl) (Sigma-Aldrich®, cat. no. L9650)
- MgCl2.6H2O (Duchefa Biochemie, cat. no. M0533)
- Midori Green Advance DNA/RNA dye (Nippon Genetics, cat. no. MG04)
- Monobasic Potassium Phosphate (KH2PO4) (Sigma-Aldrich®, cat. no. P0662)
- Murashige & Skoog medium with vitamins (Duchefa Biochemie, cat. no. M0245)
- NP-40 (Thermo Scientific, cat. no. 85125)
- Phenol (Sigma-Aldrich®, cat. no. P4557)
- Plant agar (Duchefa Biochemie, cat. no. P1001)
- PMSF Protease Inhibitor (Roche™, cat. no. PMSF-RO)
- Potassium chloride (KCl) (Duchefa Biochemie, cat. no. P0515)
- Proteinase K (Invitrogen™, cat. no. 25530049)
- Salmon sperm (Invitrogen™, cat. no. AM9680)
- Sodium chloride (NaCl) (Duchefa Biochemie, cat. no. S0520)
- Sodium desexychlae (NaDOC) (Sigma-Aldrich®, cat. no. 218928)
- Sodium dodecyl sulphate (SDS) (Sigma-Aldrich®, cat. no. 75746)
- Spermine tetrahydrochloride (Sigma-Aldrich®, cat. no. 85610)
- Streptavidine T1 Dynabeads™ MyOne™ (Invitrogen™, cat. no. 65601)
- Sucrose (Duchefa Biochemie, cat. no. S0809)
- Tris-HCl (Duchefa Biochemie, cat. no. T1501)
- Trisodium citrate (Sigma-Aldrich®, cat. no. S1804)
- Triton X-100 (Thermo Scientific Chemicals, cat. no. A16046.AP)
- Tween 20 (Millipore®, cat. no. 655206)
- β-mercaptoethanol (Sigma-Aldrich®, cat. no. M6250)

Supplementary method 2: Kits and specific materials

- Guide-it sgRNA Screening Kit (TaKaRa Bio, cat. no. 631440)
- Low Protein Binding Microcentrifuge Tubes 1.5 mL & 2 mL (Thermo Scientific, cat. no. 90410 & 88379)
- MinElute PCR Purification kit (Qiagen, cat. no. 28006)
- Platinium™ Multiplex PCR Mastermix (Applied Biosystems™, cat. no. 4464268)
- PowerUp™ SYBR™ Green Master Mix for qPCR (Thermo Scientific™, cat. no. A25777)
- PrimeScript™ RT Master Mix (Perfect Real Time) (TaKaRa Bio, cat. no. RR036A)
- QIAquick PCR Purification Kit (Qiagen, cat. no. 28104)
- QuantStudio™ 3 Real-Time PCR System (Applied Biosystems™, cat. no. A28137)
- Tissue culture boxes (Duchefa Biochemie, cat. no. S1686 or E1650)

Supplementary method 3: Solutions

***PBS***

8.0 g/L NaCl, 0.2 g/L KCl, 1.42 g/L Na_2_HPO_4_, 0.24 g/L KH_2_PO_4_.

***Home-made NEBuffer r3.1***

100 mM NaCl, 50 mM Tris-HCl (pH 8.1), 10 mM MgCl_2_, 100 µg/mL Albumin.

***10× Cas9 buffer (Ishii et al., 2019)***

200 mM HEPES (pH 7.5), 1 M KCl, 50 mM MgCl2, 50% glycerol, 10% BSA, 1% Tween 20; store at -20°C.

***Block solution (for a 400 µL aliquot) (Ishii et al., 2019)***

40 µL of 10 mM DTT (final concentration: 1 mM), 40 µL *10× Cas9 buffer* (final concentration: 1×), 320 µL ddH_2_O. Prepare fresh and keep on ice until use.

***RNP solution (for a 100 µL aliquot) (Ishii et al., 2019)***

10 µL of 10 mM DTT (final concentration: 1 mM), 10 µL *10× Cas9 buffer* (final concentration: 1×), 80 µL ddH_2_O. Prepare fresh and keep on ice until use.

***Nuclei wash buffer with Triton (Lee et al., 2017; Liu et al., 2018)***

0.1% v/v Triton X-100, 10 mM EDTA, 0.5 mM EGTA, 10 mM HEPES (pH 6.5); prepare fresh using sterile stock solutions and keep on ice until use.

***Nuclei wash buffer without Triton (Lee et al., 2017)***

200 mM NaCl, 1 mM EDTA, 0.5 mM EGTA, 10 mM HEPES (pH 6.5); filter-sterilize using a 0.45-μm filter or prepare using sterile solutions and store at RT).

***Nuclear lysis buffer (Liu et al., 2018)***

50 mM Tris-HCl (pH 8.0), 1 mM EDTA, 1 mM PMSF, 0.5% SDS, 1 mM DTT, 1 x Complete Protease cocktail inhibitor; prepare fresh using sterile stock solutions. Keep on ice until use.

For the Complete Protease cocktail inhibitor, it is possible to prepare a 25x solution by dissolving 1 pill in 2 mL ddH_2_O and filter-sterilizing with a 0.45 µm-filter and store at -20°C.

***Radioimmunoprecipitation (RIPA) 0.3 buffer (Liu et al., 2018)***

10 mM Tris·Cl (pH 7.4), 1 mM EDTA (pH 8.0), 0.3 M NaCl, 0.1% SDS, 1% v/v Triton X-100, 0.1% NaDOC, 1 mM DTT, 1 mM PMSF, 1 x Complete Protease cocktail inhibitor; prepare fresh using sterile stock solutions and keep on ice until use.

***High salt wash buffer (Liu et al., 2018)***

50 mM HEPES (pH 7.5), 1 mM EDTA (pH 8.0), 0.5 M NaCl, 1% v/v Triton X-100, 0.1% NaDOC ; store at 4°C for up to 6 months.

***LiCl wash buffer (Liu et al., 2018)***

10 mM Tris·Cl (pH 8.1), 1 mM EDTA (pH 8.0), 250 mM LiCl, 0.5% NP-40, 0.5% NaDOC; store at 4°C for up to 6 months.

***TE buffer (Liu et al., 2018)***

10 mM Tris·Cl (pH 7.5); 1 mM EDTA (pH 8.0); store at room temperature for up to 6 months.

***SDS elution buffer (Liu et al., 2018)***

50 mM Tris·HCl (pH 8.1); 10 mM EDTA (pH 8.0), 1% SDS; Store at room temperature for up to 6 months.

***20X SSC***

3 M NaCl, 300 mM trisodium citrate. Adjust pH to 7.0 with HCl and store at 4°C.

Supplementary method 4: Methods

In vitro amplification of the fragment containing the telomeric repeat

We designed 2 pairs of primers to amplify 2 fragments, (short region, SHRT_Fw and SHRT_Rv) and 400 bp (long region, LONG_Fw and LONG_Rv) in length, each of them containing a telomeric spacer and a PAM at one of its extremities. (**Figure 2A**) The content of the fragment is of no importance as long as they do not contain another telomeric fragment. For each pair of primers, one primer (LONG_Fw and SHRT_Rv) aligns only with the template sequence for amplification and is ∼20 b long while the other one aligns with the target sequence for >20b and carries an extension with the telomeric spacer and the PAM (SHRT_Fw) or in the reverse direction (LONG_Rv). Then, the final fragment is generated as follows.

- Amplify separate reactions the two sub-fragments by PCR using Platinium™ Multiplex PCR Mastermix (Applied Biosystems™, cat. no. 4464268), two 50 mL reactions at the same time (35 cycles composed of 94°C for 30 sec, 58°C for 30 sec, 72°C for 30 sec). In case the difference between the TM of the two primers of a pair is too different, it is possible to do 5 cycles with primer pairing at 68°C and 30 cycles with pairing at 58°C. Other parameters remain unchanged.
- Run both reactions on a 2 % agarose gel for 35 min at 150 V, cut the bands correspond to the desired lengths and purify them using Qiaquick gel purification kit.
- Assemble the two sub-fragments in 4-8 50 mL PCR reactions containing the two primers without spacer and PAM (LONG_Fw and SHRT_Rv), 20 ng of each sub-fragments per reaction (28-35 cycles composed of 95°C for 10 sec; 58°C for 15 sec; 68°C for 20 sec).
- Purify the PCR product with Qiaquick PCR purification kit and run on 2% agarose gel for 35 minutes to assess the integrity of the fragment.

We also amplified a fragment containing a unique spacer for the bacterial GAL4 gene (Liu *et al.*, 2018) followed by a PAM of the NGG sequence. Note that only the SHRT_Fw and LONG_Rv primers are spacer specific; the two other primers are usable in any case.

Quick explanation of the principle of the CRISPR-FISH technique.

The CRISPR-FISH technique (Potlapalli *et al.*, 2024) uses a catalytically inactive dCas9 complexed with a fluorescently labeled guide RNA (gRNA) composed of 2 put-together parts: the CRISPR-RNA (crRNA) that carries the spacer sequence, and a trans-activating-RNA (tracrRNA) that partially hybridizes with the first one and insures the stability of the gRNA and the specificity of the binding of the dCas9. This complex binds to the target DNA sequence in fixed, isolated nuclei from fresh plant leaves. Fluorescence microscopy is used to detect the bound complexes, revealing the spatial distribution of the target loci (Dreissig *et al.*, 2017; Ishii *et al.*, 2019; Němečková *et al.*, 2019; Potlapalli *et al.*, 2023). It is one of the new imaging techniques that emerged with the non-editing use of CRISPR-Cas9 system (Devillars *et al.*, 2024). This approach avoids DNA denaturation, making it gentler and more compatible with immunostaining and chromatin-sensitive assays. For the purpose of our experiments, we have been using a biotinylated dCas9 loaded with a gRNA with the telomeric spacer (Potlapalli *et al.*, 2020) or, for negative controls, either the gRNA with spacer for Gal4 (Liu *et al.*, 2018) or only the tracrRNA part of the gRNA.

Quantitative analysis of CRISPR-FISH and FISH foci

The procedure uses ImageJ/Fiji is given for either all FISH images or all CRISPR-FISH images. It is recommended proceed with all images at once using a Fiji macro with iteration cycles.

- For all FISH or CRISPR-FISH images, open the DAPI channel and apply automatic threshold using Triangle method with *dark* parameter to exclude low-intensity background.
- Run Convert to Mask, Close- and Erode to delimitate the nuclei.
- Run Watershed if the delimitation is not clear enough.
- Apply Analyze Particle with *size=100-Infinity pixel exclude add* parameters to and save them as ROI.
- Open FISH or CRISPR-FISH channels of all images.
- Apply Top Hat with *radius=3* for CRISPR-FISH or *radius=20* for FISH

Now, proceed as follows for each nucleus taken independently

- Duplicate the FISH or CRISPR-FISH channel and remove everything but the given nuclei with Clear Outside
- Apply MaxEntropy auto threshold with *dark* parameter
- Run Convert to Mask to delimitate the foci
- Run Watershed if the delimitation is not clear enough.
- Apply Analyze Particles with *size=N-Infinity pixel display exclude summarize add*, where *N=3* for CRISPR-FISH foci, *N=1* for FISH.
- The number of foci and their characteristics are visible in the Measure table.
- To limit false discovery due to a threshold issue, disregard all nuclei showing a mean signal size of less than 4 pixels.

Alignment of FISH and CRISPR-FISH channels to produce the colocalization images.

- Open the channels of the two images of one field in GIMP 2.10.34 as layers in different layer groups (one group for the CRISPR-FISH and its DAPI, one group of the FISH and its DAPI). Hide both non-DAPI channels.
- Move one whole layer group so that the DAPI of both groups are superposed.
- Select the rectangle corresponding to the intersection of the layers and crop the image according to the selection, delete one of the DAPI layers.
- In the layer window, move all the layers out of the layer groups and delete the empty layer groups.
- Export the resulting image as a tiff file that will be multipage.
- Open this tiff file with ImageJ and split the stack back into single images.
- You can now produce the merged image by merging the images.

Cleavage assay with Cas9 and gRNAs.

We used the reagents of the Guide-it sgRNA Screening Kit and performed as follows:

- For each condition, combine in a 200-μl PCR tube 1 µL of assembled (if needed) gRNA or control RNA (50 ng/µL) and 0.5 µL of Guide-it Recombinant Cas9 Nuclease (500 ng/μL). If you have multiple samples, make a master mix. Incubate for 5 min at 37°C in a thermocycler.
- Add to each PCR-tubes 100 ng of the amplified fragment containing the spacer specific for this gRNA followed by a PAM or control fragment, 1µL of 15X Cas9 Reaction Buffer, 1 µL of 15X BSA and 6.5 µL RNase Free Water.
- Mix well and very gently by pipetting. Incubate using a thermal cycler with the following conditions: 37°C for 1h, 80°C for 5 min and 4°C forever.
- Run the entire sample on a 2% agarose gel along with an appropriate DNA marker as the control and telomeric fragments alone.

Binding assay

To evaluate the specificity and efficiency of the ribonucleoprotein complex, consisting of the gRNA and dCas9, in recognizing and binding to the target telomeric sequences, perform a binding assay as described by (Zou *et al.*, 2021). In our case, to increase the discriminative propriety of the gel, the length fragment for this experiment was reduced to 391 bp by PCR amplification using internal primers (amp_fram_Fw and amp_fram_Rv). After controlling the size and concentration of DNA amplicon, the PCR mix can be used as template with no purification required.

- Mix the following components in PCR tubes, in order: 8.1 µL Nuclease free water, 1 µL home-made NEBuffer r3.1, 0.5 µL of 10 µM fragment-specific or unspecific gRNA, 0.4 µL of 10 µM dCas9.
- Leave at room temperature for 30 min.
- Add 100 ng of amplified telomeric fragment.
- Add 0.5 µL of 20 µg/µL Proteinase K in one of the tubes with specific gRNA
- Incubate in a thermocycler at 55°C for 15 minutes, then 4°C for infinite time.
- Run on 2% agarose gel stained with 0.1% Midori Green Advance for 30 min.

Nuclei extraction from plant leaves for the nuclei transfection assay

- Freeze the leaves (up to 1g) in liquid nitrogen and grind them with mortar and pestle. If needed, grind in different batches.
- Immerse the leaf powder in 5 mL LB01 buffer in a 50-mL falcon and mix by shaking the tube several times.
- Pre-filter the lysate through a 60 µm mesh filter then filter it through a 30 µm mesh cytology filter and collect the flow-through in 15 mL falcon.
- Centrifuge at 3 000 × g for 10 min, 4°C and discard the supernatant.
- Resuspend the pellet in 2 mL ice-cold PBS, centrifuge at 3 000 × g for 2 min, 4°C and discard the supernatant.
- Resuspend the pellet in 1-2 mL ice-cold PBS.
- Load 10 µL of a ≥ 20x dilution on the Denovix CellDrop and count the nuclei using the Brightfiled program and the following parameters: Chamber Height: 100; DirectPipette (or Slide): Pipette; Dilution Factor: your dilution; Protocol Min Diameter: 2 µm; Protocol Max Diameter: 20 µm; Roundness: 20; Small Cell Mode: False; Irregular Cell Mode: False. Interpret the values of “Cell Count” as “Nuclei Count” and “Cell/mL” as “Nuclei/mL”.

Nuclei transfection assay

- Prepare RNP complex as described in the next sections, adding 4 µL of dCas9 and 4 µL of specific or unspecific gRNA with ATTO-550 fluorophore instead of 1 µL each.
- Isolate nuclei from plant leaves as described previously.
- Pellet the desired number of nuclei at 3 000 × g for 2 min and resuspend them in 400 µL PBS.
- Add 100 µL of RNP to the nuclei and incubate at 37°C for 2 hours on a wheel.
- Add 13.88 μL of 37% formaldehyde (1% FA)
- Let the tube invert on the wheel for 15 min, RT.
- Add 102.8 µL of 600 mM glycine (0.1 M glycine)
- Let the tube invert on the wheel for 5 min, RT.
- Centrifuge at 3 000 × g for 2 min.
- Resuspend the pellet in 500 µL PBS.
- Load 10 µL of a ≥ 20x dilution on the Denovix CellDrop and count the nuclei using the IP program and the following parameters: Chamber Height: 100; DirectPipette (or Slide): Pipette; Dilution Factor: your dilution; Protocol Min Diameter: 2 µm; Protocol Max Diameter: 20 µm; Live Roundness: 20; Dead Roundness: 20; Red Fluorescence Threshold: 20 Small Cell Mode: False; Irregular Cell Mode: False. Interpret the values of “% Viability” as “% of non-transfection”, “Live Cell/mL” as “Non-transfected nuclei/mL”, “Dead Cell/mL” as “Transfected nuclei/mL” and “Total Cell/mL” as “Total Nuclei/mL”.

Design of the spacers and primers for low-repetition or single loci

- Retrieve the sequence of all genes (here STS genes) in the reference genome.
- Analyze them for occurrences of 23mers with Jellyfish 2.3.0.
- Discard the 23mers that do not either start with CCN or finish with NGG (i.e. non constituting a valid spacer + PAM).
- Blast the remaining 23mers back on all query sequences, outputting the results as tabular file(s) indicating, for each sequence, the exact start and end positions of all 23mers completely aligned on the queries.
- These tables can be imported in R, along with the genome annotation files, localize the repetitions of the spacers in the sequences. Select few spacers based on your experimental interests.
- Control the on-score of each selected spacer with CRISPR-P version 2.0. (http://crispr.hzau.edu.cn/CRISPR2/help.php) or Geneious Prime version 2025.0.
- Use Geneious Prime to design primers suited for qPCR on precipitated and input genomic DNA. The amplified fragment should be ~150 bp and totally comprised in a -300 to +300 pb-window from the spacer.
- Design and order the primers to amplify the synthetic fragment for cleavage assay, that should be identical to those for telomeres and Gal4, except for parts corresponding to spacer and PAM.
- Design and order the primers to amplify the fragments for T7-mediated RNA synthesis.
  - PCR-amplify a first intermediary fragment from 5 ng of the AtU6 vector using 2 µL of the gRT-Fw primer that anneals at the beginning of the gRNA scaffold and contains an extension consisting in the 20 nt of a given spacer, 2 µL of gR-R primer that anneals 13 nt downstream of the scaffold’s end (Ma *et al.*, 2015), 50 µL Platinium PCR Master Mix and 5 µL GC enhancer, for a total of 100 µL PCR mix. Split in two 50 µL-aliquots in PCR tubes and run the PCR. Use the following PCR parameters: 30 sec at 94°C then 30 sec at 94°C and 35 cycles composed of 94°C for 30 sec, 58°C for 30 sec, 72°C for 10 sec, then final 7 min of extension at 72°C. Purify the fragment of DNA with QIAquick PCR Purification kit.
  - Amplify the DNA fragment serving as template for T7 *in vitro* transcription by performing PCR on 5 ng of the gRT fragment, 1 µL T7-Fw primer (annealing on a given spacer and containing the T7-promoter as extension), 1 µL of Scaffold_Rv primer (same for all spacers and aligning with the scaffold-part of the gRNA encoding sequence), 25 µL Platinium PCR Master Mix, 2.5 µL GC enhancer, and RNase-free water up until 50 µL. To design the promoter primer, refer to the guidelines of the Guide-it sgRNA Screening Kit. Use the following PCR parameters: 30 sec at 94°C and 35 cycles composed of 94°C for 30 sec, 68°C for 30 sec, 72°C for 9 sec, then final 7 min of extension at 72°C.
  - Retrotranscribe DNA template and subsequently purify the obtained DNA template using the appropriate section of the Guide it gRNA sgRNA synthesis-kit. Quantify the gRNA concentration with Nanodrop.
- To test the workability of the gRNA, perform cleavage as described before using the Guide it gRNA sgRNA synthesis-kit. Store the synthetized gRNAs at -20°C.
- Once positive result for the cleavage experiment is obtained, the gRNA can be used for precipitation. Dilute it to 10 µM (or 303 ng/µL for a 103 nucleotide-long single-stranded RNA) in RNase-free water and store at -20°C.

Supplementary method 5: TROUBLESHOOTING points

Counting of the telomeric repeats in grapevine T2T genomes

A small number of telomeric repeats and counted telomeric kmers may come from an incorrect assembly of those in the chosen genome. Therefore, it is important to use Telomere2Telomere (T2T) assemblies, that contain all chromosomes from one extremity to the other (at least for the large majority) and few unplaced sequences. Indeed, when using a non-T2T assembly, the telomeres, as highly repetitive sequences, are very likely to be missing.

Microscopy

- As already mentioned in the manuscript, the feasibility of CRISPR-FISH depends on the number of telomeric repeats. If transferring the protocol to another species that the ones already tested in literature, be sure to check whether the telomeres are long enough to.
- If the CRISPR-FISH foci are not very clear, then it may come from an unproper post-fixation of the sample on the slide. It is recommended to start again or to adjust post-fixation procedure, but it may still be possible to produce exploitable images by performing Z-stack captures.
- Also, be aware to maintain the RNP solution and the slides in the dark to avoid bleaching of the ATTO fluorophore on the tracrRNA.

Precipitation protocol – Nuclei isolation

After having isolated nuclei from plant leaves and before going on with the transfection phase, it is of good usage to control the number of nuclei with the Cell Counter. We do not recommend to start the process with an amount lower than 10^7^ nuclei per transfection. Going below this limit may lead to an insufficient quantity of DNA material in the end.

Precipitation protocol – Sonication

- Sonication is known for being species- and tissue- specific. Whenever applying the protocol to another plant or another part of the plant, it is important to verify that the chromatin is sheared correctly. Inadequate sonication will lead to either too long or too short chromatin fragments, which can compromise the efficiency of the precipitation and/or NGS sequencing.
- At the end of the sonication part of the protocol, it is important to collect enough Input sample. Therefore, it is possible to first save 20 µL of the supernatant form the upper part of the tube without disturbing the pellet, and then to carefully take of the rest of the supernatant (approximately 400 µL).

Precipitation protocol – Chromatin precipitation with streptavidin magnetic beads

- When collecting and washing the Dynabeads, especially if you have a lot of samples, thoughtfully resuspend the beads vial every 2 pipetting otherwise they will fall in the bottom. It is possible either to wash the beads from all samples together and split them after the last wash or to wash the beads for each sample on their own. The first option will often lead to a different number of beads into the tubes, since the beads easily attach to the pipette tip, and is to consider only for a limited number of samples. The second option can lead to having to prepare and consume a high quantity of RIPA buffer. In that case, since the quantity of beads for a single sample is very small, it is possible to wash them in lower amount of RIPA buffer, such as 500 µL.
- During the washes with SDS, LiCl Wash buffer, and High Salt Wash Buffer, It is preferable not to use filter tips since they will produce a high quantity of bubbles which will make the pipetting difficult and even more when using a filter.
