## Supplementary material for "GRASP: A PLANT TRANSFORMATION-INDEPENDENT CRISPR-BASED SYSTEM FOR AFFINITY PURIFICATION OF SPECIFIC CHROMATIN LOCI": Supplemetary File 1

BASHSCRIPT: Jellyfish_blast_Rgraphs_script

```
module load blast+
module load jellyfish
module load samtools
if [ $
then
read -p "Do JELLYFISH [y/n] : "  JELLY
read -p "Do BLAST [y/n] : "  BLAST
read -p "Do GRAPH [y/n] : "  GRAPH
read -p "Which species (enter one ALL IN MINUCULES) : " SPECIES
else
JELLY=$1 ;  BLAST=$2 ; GRAPH=$3 ; SPECIES=$4
fi
GENERAL_OUTDIR= ### # Your output directory
GENOME_FILE= ### #Where to find the genome fasta file
GENOME_DIRECTORY=$(dirname $GENOME_FILE)
EXT=$(basename $GENOME_FILE | rev | cut -d '.' -f 1 | rev)
GENOME_NAME=$(basename --suffix=.$EXT $GENOME_FILE)
OUTDIR_JELLYFISH=$GENERAL_OUTDIR/jellyfish_kmeri_telomeri/$SPECIES
OUTDIR_BLASTDB=$GENOME_DIRECTORY/${GENOME_NAME}_blastdb
DB=$OUTDIR_BLASTDB/$GENOME_NAME
OUTDIR_BLASTN=$OUTDIR_JELLYFISH/BLAST_out_telomeres
kmers_telomers=$GENERAL_OUTDIR/list_kmers_telomeres.txt
tabular_out=$OUTDIR_JELLYFISH/BLAST_tabular_out_$SPECIES.txt
echo Species:$SPECIES, Genome:$GENOME_NAME
mkdir -p $OUTDIR_JELLYFISH
if test $JELLY == "y"
then
    JF_FILENAME=$OUTDIR_JELLYFISH/$GENOME_NAME
    [[ ! -f ${JF_FILENAME}.jf ]] && jellyfish count -m 21 -s 100M -t 10 -C  -o ${JF_FILENAME}.jf $GENOME_FILE
    echo done jellyfish Count
	jellyfish dump -c -L 100 ${JF_FILENAME}.jf > ${JF_FILENAME}.txt && rm ${JF_FILENAME}.jf
	grep -f $kmers_telomers ${JF_FILENAME}.txt
    echo done jellyfish dump
	echo done jellyfish
fi
if test $BLAST == "y"
then
    mkdir -p $OUTDIR_BLASTDB
    makeblastdb -dbtype "nucl" -parse_seqids -blastdb_version 5 -taxid 29760 -in $GENOME_FILE -out $OUTDIR_BLASTDB/$GENOME_NAME
    mkdir -p $OUTDIR_BLASTN
    compteur=0
    total=$(grep -c -f $kmers_telomers ${JF_FILENAME}.txt)
    for QUERY in $(grep -o -f $kmers_telomers ${JF_FILENAME}.txt)
    do
        TEMPFILE=$OUTDIR_JELLYFISH/temp_queryfile_$QUERY.fa
		echo ">${QUERY}" >> $TEMPFILE
		echo $QUERY >> $TEMPFILE
        blastn -task blastn-short -db $DB -query $TEMPFILE -out $OUTDIR_BLASTN/${QUERY}.txt -outfmt 6  -perc_identity 100
        ((compteur++))
        echo $HAP $QUERY  $compteur \/ $total
    if [ ! -s $OUTDIR_BLASTN/${QUERY}.txt ]
    then
        echo Empty file
        rm $OUTDIR_BLASTN/${QUERY}.txt
    fi
    done && rm ${JF_FILENAME}.txt && echo done blast
fi
[[ -f $GENOME_FILE ]] && samtools faidx $GENOME_FILE
mv $GENOME_FILE.fai $OUTDIR_JELLYFISH
cat $OUTDIR_BLASTN/* > $tabular_out
wc -l $tabular_out
if test $GRAPH == "y" -a -s $tabular_out
then
SCRIPT_FILE=$GENERAL_OUTDIR/chr_k21_telomeric_GENERAL.r
R_FILE=$GENERAL_OUTDIR/chr_k21_telomeric_GENERAL_$SPECIES.r
fai_file=$OUTDIR_JELLYFISH/$GENOME_NAME.$EXT.fai
START_VARIABLES="OUTDIR = \"$OUTDIR_JELLYFISH\" ; \
fai.file = \"$fai_file\" ; \
blast.file = \"$tabular_out\" ; \
SPECIES = \"$SPECIES\""
mkdir -p $(dirname $R_FILE)
echo -e $START_VARIABLES > $R_FILE
cat $SCRIPT_FILE >> $R_FILE
Rscript $R_FILE
fi
```

RSCRIPT: chr_k21_telomeric_GENERAL

```
library(data.table)
library(ggplot2)
SPECIES = "Sultana Génome T2T"
OUTDIR = file.path(".....",SPECIES) #YOUR FOLDER
fai.file = grep(".fai", file.path(OUTDIR,dir(OUTDIR)),value = TRUE)
blast.file = grep("rm_superposed" ,grep("BLAST_tabular_out_", paste0(OUTDIR,"/",dir(OUTDIR)),invert = F,value = TRUE),invert = TRUE,value = TRUE)
KMER_LENGTH = 21
if(SPECIES %in% c("Sultana Génome T2T","thompson_seedless_t2t")) {CHROMOSOME_NAMES_PATTERN <- "Chr" ; CUSTOM_REPLACEMENT <- NULL ; LAT_NAME = "Vitis vinifera cv Thompson Seedless"}
if(SPECIES %in% c("Pinot Noir Génome T2T","pinot_noir_t2t","pinot_noir_alt_t2t")) {CHROMOSOME_NAMES_PATTERN <- "chr" ; CUSTOM_REPLACEMENT <- NULL ; LAT_NAME = "Vitis vinifera cv Pinot Noir"}
if(SPECIES %in% c("Maïs Gatersleben","maize")) {CHROMOSOME_NAMES_PATTERN <- "CM0" ; CUSTOM_REPLACEMENT <- function(chrom) as.integer(gsub(CHROMOSOME_NAMES_PATTERN,"",chrom))-39149 ; LAT_NAME = "Zea mays"}
if(grepl("arabidopsis", SPECIES,ignore.case = TRUE)) {CHROMOSOME_NAMES_PATTERN <- "chr" ; CUSTOM_REPLACEMENT <- NULL ; LAT_NAME = "Arabidopsis thaliana"}
if(grepl("cabernet_franc_hap" , SPECIES)) {CHROMOSOME_NAMES_PATTERN <- "VITVvi_vCabFran04_v1.1.hap..chr" ; CUSTOM_REPLACEMENT <- NULL ; LAT_NAME = "Vitis vinifera cv. Cabernet Franc"}
if(grepl("fennel_long_reads" , SPECIES)) {CHROMOSOME_NAMES_PATTERN <- "LG" ; CUSTOM_REPLACEMENT <- function(x) sapply(x, function(y) match(y, reps$chr)) ; LAT_NAME = "Foeniculum vulgare" }
if(grepl("fennel_long_reads" , SPECIES)) {CHROMOSOME_NAMES_PATTERN <- "LG" ; CUSTOM_REPLACEMENT <- function(x) sapply(x, function(y) match(y, reps$chr)) ; LAT_NAME = "Foeniculum vulgare" }
MAX_SECTOR_NUMBER = 50
THRESHOLD_KMER_PRINT  = 10
THRESHOLD_KMER_RETAIN = 20
ESPACE_X_CHROMOSOME = 0.45
ESPACE_Y_REPS = 1/200
MAX_ITERATION = 2
LOOP_KMER = F
KMER_UNO = T
if (LOOP_KMER) KMER_UNO = c(TRUE,FALSE)
STAMPA_PDF = F
STAMPA_TAB = F
STAMPA_SVG = T
COULR_MIN = "transparent"
COULR_MAX = "#B10E0C"
COULR_SECOND = "black"
last<-function(x) {return(x[length(x)])}
numbers_between <- function(lst, lower, upper) {
  return(which(lst > lower & lst < upper))
}
is_between <- function(number, lower_bound, upper_bound) {
  return(number >= as.numeric(lower_bound) & number <= as.numeric(upper_bound))
}
change_chr_names = function(x) {
  if (iteration == 0) {
    if (!is.null(CUSTOM_REPLACEMENT)) return(CUSTOM_REPLACEMENT(x))
    return(as.integer(gsub(pattern = CHROMOSOME_NAMES_PATTERN, replacement = "",x)))
  } else {
    all_chr = unique(x)
      return(sapply(x,function(y) {which(y == all_chr)}))
    }
  }
convertir_unite <- function(nombre, R = 0) {
  if (nombre >= 1e9) {
    return(paste0(round(nombre / 1e9, R), " Gbp"))
  } else if (nombre >= 1e6) {
    return(paste0(round(nombre / 1e6, R), " Mbp"))
  } else if (nombre >= 1e3) {
    return(paste0(round(nombre / 1e3, R), " kbp"))
  } else {
    return(paste0(round(nombre, R), " bp"))
  }
}
progressing_bar = function(it, tot, series = NULL) {
  if (is.null((series))) {
    if (it == 1) return(" (0%) [|")
    if (it %in% round(tot/2:9,0) ) return("|")
    if (it == tot) return("|] (100%) \n")
    return()} else {
      if (it == series[1]) return(paste0(" (1/",length(series),") [="))
      if (it == series[length(series)]) return(paste0(">] (",length(series),"/",length(series),")\n"))
      return("=")
    }
}
raw_chr = function() {
  fai.table <- fread(fai.file, data.table=F, header=F,col.names = c("chr","length", "start",NA,NA))[,1:3]
  fai.table$length = as.integer(fai.table$length)
  fai.table$end = as.numeric(fai.table$start) + as.numeric(fai.table$length)
  fai.table$side = fai.table$chr
	return(fai.table)
  }
telomeric_chr = function(k_sect_tab = kmer_sector_tab) {
  telomeric_sectors = data.frame(
    kmer = character(0),
    chr = character(0),
    length = numeric(0),
    start = numeric(0))
    cat("\n\n Computing zoom on telomeric regions...\n")
    for (kmer in kmer_list) {
	  cat(">",kmer)
    in_sector = FALSE
    for (i in 1:nrow(k_sect_tab[[kmer]])){
      cat(progressing_bar(i,nrow(k_sect_tab[[kmer]])))
      if (k_sect_tab[[kmer]]$Repetitions[i] >= THRESHOLD_KMER_RETAIN & in_sector == FALSE & !is.na(k_sect_tab[[kmer]]$Repetitions[i])) {
        in_sector = TRUE
        chm = k_sect_tab[[kmer]]$chr[i]
        deb = k_sect_tab[[kmer]]$start_sector[i]
      }
      leave_side = eval(i == nrow(k_sect_tab[[kmer]]) | k_sect_tab[[kmer]]$side[i] != k_sect_tab[[kmer]]$side[i+1])
      if ((leave_side == TRUE | k_sect_tab[[kmer]]$Repetitions[i+1] < THRESHOLD_KMER_RETAIN  ) & in_sector == TRUE) {
        in_sector = FALSE
        len = k_sect_tab[[kmer]]$end_sector[i] - deb
        telomeric_sectors[nrow(telomeric_sectors) + 1,] = c(kmer,
                                                            chm,
                                                            len,
                                                            deb + chr_dim$start[chr_dim$side == k_sect_tab[[kmer]]$side[i]])
      }
    }
  }
  telomeric_sectors = unique(telomeric_sectors[,c("chr","length","start")])
  dupls = unlist(sapply(unique(telomeric_sectors$start), function(el) {
    if (length(subset(telomeric_sectors, start == el)$start) != 1) {
      enleve = min(subset(telomeric_sectors, start == el)$length)
      return(which(telomeric_sectors$start == el & telomeric_sectors$length == enleve))
    }
  }))
  if (length(dupls) > 0) telomeric_sectors = telomeric_sectors[-dupls,]
  numeros = data.frame()
  for (chrom in unique(telomeric_sectors$chr)) {
    e = subset(telomeric_sectors, chr == chrom)[,c("chr","start")]
    e$side = paste0(e$chr,"-",1:nrow(e))
    numeros=rbind(numeros,e)
  }
  telomeric_sectors = merge(telomeric_sectors, numeros, by = c("chr","start"))
  telomeric_sectors$length = as.numeric(telomeric_sectors$length)
  telomeric_sectors$start = as.numeric(telomeric_sectors$start)
  telomeric_sectors$end = telomeric_sectors$start + telomeric_sectors$length
    return(telomeric_sectors[order(as.numeric(telomeric_sectors$start)),])
}
raw_tab = function() {
  rmsup.blast.file = paste0(dirname(blast.file),"/rm_superposed_kmers_",basename(blast.file), collapse = "")
  blast_zero_name = paste0('blast.table_zero_',SPECIES, sep = "")
  if ( ! exists(blast_zero_name)) { blast.table<-subset(read.table(blast.file, header=F, col.names = c("kmer","chr",NA,"align",rep(NA,4),"pos",rep(NA,3))),
                      ! chr %in% c("Vv_mitochondrion","Vv_chloroplast") & align == KMER_LENGTH)
  row.names(blast.table) = 1:nrow(blast.table)
  if (! basename(rmsup.blast.file) %in% dir(dirname(blast.file))) {
    cat("\n Recherche des kmers superposés: ", rmsup.blast.file,"\n")
    blast.table$nn = 1:nrow(blast.table)
    sup = integer(0)
    for (K in 1:nrow(blast.table)) if ( ! K %in% sup ) sup = c(sup, subset(blast.table, kmer == blast.table$kmer[K] & chr == blast.table$chr[K])[
      numbers_between(blast.table$pos[blast.table$kmer == blast.table$kmer[K] & blast.table$chr == blast.table$chr[K]],
                      blast.table$pos[K],
                      blast.table$pos[K]+KMER_LENGTH-1), "nn"] )
    write.table(x = sup, file = rmsup.blast.file, col.names = F, row.names = F) }
  else {
    cat("\n Fichier des kmers superprosés: ", rmsup.blast.file,"\n")
    sup = read.table(rmsup.blast.file, header = F)[,1]
      }
  blast.table = blast.table[-sup,c("kmer","chr","pos")]
  assign(blast_zero_name, blast.table, envir = .GlobalEnv)
  } else {
    assign("blast.table", get(blast_zero_name))
  }
  blast.table$posdecale <- as.numeric(blast.table$pos) +
    sapply(blast.table$chr, function(chrom) {
      index <- match(chrom, chr_dim$chr)
        return(chr_dim$start[index])
    })
  blast.table$side = blast.table$chr
    return(blast.table[grep(invert = TRUE, "Query_1", blast.table$kmer),])
}
telomeric_tab = function() {
  k_tab$side <<- NA
  for (i in 1:nrow(chr_dim)) {
    deb =  chr_dim$start[i]
    end = chr_dim$start[i] + chr_dim$length[i]
    chrom = chr_dim$chr[i]
    in_chr = intersect(numbers_between(k_tab$posdecale,deb, end),which(k_tab$chr == chr_dim$chr[i]))
    k_tab$side[in_chr] <<- chr_dim$side[i]
}
  k_tab <<- subset(k_tab, !is.na(side))
}
calc_xy = function(dim_tab, by = side){
dim_tab = subset(dim_tab, side %in% reps$side)
rownames(dim_tab) = 1:nrow(dim_tab)
dim_tab$"chr_num" = change_chr_names(dim_tab$side)
dim_tab <- subset(dim_tab, chr_num != 0)
dim_tab$"xmin"<-dim_tab$"chr_num"-ESPACE_X_CHROMOSOME
dim_tab$"xmax"<-dim_tab$"chr_num"+ESPACE_X_CHROMOSOME
return(dim_tab)
}
calc_sectors = function() {
  start_sector_all<-NULL
  end_sector_all<-NULL
  chr_sector_all<-NULL
  side_sector_all<-NULL
for (i in 1:nrow(chr_dim)) {
  chr=chr_dim$"chr"[i]
  side=chr_dim$"side"[i]
  numsector<-chr_dim$sectors[i]
  lengthsector<-round(chr_dim$"length"[i]/numsector, 0)
  start_sector<-c(0,lengthsector+1)
  end_sector<-lengthsector
  while (chr_dim$"length"[i]-last(start_sector)>=lengthsector) {
    end_sector<-c(end_sector,last(start_sector)+lengthsector)
    start_sector<-c(start_sector,last(end_sector)+1)}
  end_sector<-c(end_sector,chr_dim$"length"[i])
  start_sector_all<-c(start_sector_all,start_sector)
  end_sector_all<-c(end_sector_all,end_sector)
  chr_sector_all<-c(chr_sector_all,rep(chr,length(start_sector)))
  side_sector_all<-c(side_sector_all,rep(side,length(start_sector)))
  }
  sector_all<-data.frame(side=side_sector_all,chr=chr_sector_all, start_sector=start_sector_all, end_sector=end_sector_all,
                         sect = unlist(sapply(unique(side_sector_all), function(i) 1:length(which(side_sector_all == i)))))
  sector_all$ylabel = (sector_all$"start_sector" + sector_all$"end_sector")/2
  sector_all[,c("start_sector_graph","end_sector_graph")] = sector_all[,c("start_sector","end_sector")]
  if (iteration == 0) return(sector_all)
  center_chrom = chr_dim[chr_dim$sectors == max(chr_dim$sectors),c("side","sectors")][1,]
  abs_center_sector = subset(sector_all, side == center_chrom$side)[round(center_chrom$sectors/2,0),"ylabel"]
  all_centers = unlist(sapply(unique(sector_all$side), function(chromo)
    subset(sector_all, side == chromo)[round(subset(chr_dim, side == chromo)$sectors/2,0),c("side","ylabel")]
    ))
  all_centers = data.frame(side = grep(CHROMOSOME_NAMES_PATTERN,all_centers,value = TRUE),
                    center = as.numeric(grep(CHROMOSOME_NAMES_PATTERN,all_centers,value = TRUE,invert = TRUE)))
  all_centers$diffs = abs_center_sector - all_centers$center
  sector_all = merge(sector_all, all_centers, by = "side")
  sector_all[,c("ylabel","start_sector_graph","end_sector_graph")] = sector_all[,c("ylabel","start_sector","end_sector")] + sector_all$diffs
	return(sector_all)
}
calc_kmer_sectors = function(kmers){
  cat("\n Calculating sectors...\n")
  kmer_chr_dim = list()
  kmer_sector_tab = list()
  kmer_this_chr = list()
  for (kmer in unique(kmers)) {
    cat(">",kmer)
    telomeric_kmertab<-k_tab[k_tab$"kmer"==kmer,]
    kmer_chr_dim[[kmer]]<-chr_dim
    if(iteration == 0) rep_lab_pos = 2 else rep_lab_pos = 4
    kmer_chr_dim[[kmer]]$rep_label<-sapply(chr_dim$side, function(x) max(subset(sector_tab, side == x)$end_sector_graph))+rep_lab_pos*esp_resp
    kmer_chr_dim[[kmer]]$Repetitions_this_chr<-0
    kmer_chr_dim[[kmer]]$ymin <- sapply(chr_dim$side, function(x) min(subset(sector_tab, side == x)$start_sector_graph))
    kmer_chr_dim[[kmer]]$ymax <- sapply(chr_dim$side, function(x) max(subset(sector_tab, side == x)$end_sector_graph))
    kmer_sector_tab[[kmer]]<-sector_tab
    kmer_sector_tab[[kmer]]$"Repetitions"<-0
    b = list()
    telomeric_kmertab$num_kmer = 1:nrow(telomeric_kmertab)
    g = 1
    for (chr in kmer_chr_dim[[kmer]]$side) {
      cat(progressing_bar(chr,series = kmer_chr_dim[[kmer]]$side))
      kmer_chr_dim[[kmer]]$"Repetitions_this_chr"[kmer_chr_dim[[kmer]]$"side"==chr]<-sum(telomeric_kmertab$"side"==chr)
      Repetitions<-telomeric_kmertab[telomeric_kmertab$"side"==chr,]
      for (j in grep(chr,kmer_sector_tab[[kmer]]$"side")) {
        kmer_sector_tab[[kmer]]$"Repetitions"[j]<-sum(Repetitions$"posdecale">=kmer_sector_tab[[kmer]]$"start_sector"[j]+chr_dim$start[chr_dim$side == chr] & Repetitions$"posdecale"<=kmer_sector_tab[[kmer]]$"end_sector"[j]+chr_dim$start[chr_dim$side == chr])
        k = which(Repetitions$"posdecale">=kmer_sector_tab[[kmer]]$"start_sector"[j]+chr_dim$start[chr_dim$side == chr] & Repetitions$"posdecale"<=kmer_sector_tab[[kmer]]$"end_sector"[j]+chr_dim$start[chr_dim$side == chr])
        b[[g]]  = Repetitions[k[!is.na(k)],"num_kmer"]
        g = g + 1
        }
    }
    reps[,kmer] <<- kmer_chr_dim[[kmer]]$"Repetitions_this_chr"
    sector_tab[,kmer] <<- kmer_sector_tab[[kmer]]$"Repetitions"
    kmer_this_chr[[kmer]] = b
    }
  for (liste in c("kmer_chr_dim","kmer_sector_tab","kmer_this_chr")) assign(liste, get(liste), envir = .GlobalEnv)
}
calc_graph_adds <- function(dim_tab) {
  dim_tab$ymin <- sapply(dim_tab$side, function(x) min(subset(sector_tab, side == x)$start_sector_graph))
  dim_tab$ymax <- sapply(dim_tab$side, function(x) max(subset(sector_tab, side == x)$end_sector_graph))
  dim_tab$slot_low_lim <- dim_tab$"ymin" - 2*esp_resp
  dim_tab$slot_up_lim <- dim_tab$"ymax" + 2*esp_resp
  dim_tab$slot_chr_name <- dim_tab$"ymin" - 4*esp_resp
  dim_tab$first_kmer = sapply(dim_tab$side,function(chrom) min(subset(k_tab, side == chrom)$pos))
  dim_tab$last_kmer = sapply(dim_tab$side,function(chrom) max(subset(k_tab, side == chrom)$pos))
```

dim_tab**$**start_chr **=** sapply**(**1**:**nrow**(**dim_tab**)**, **function(**i**)** **if** **(**dim_tab**$**start**[**i**]** **-** sector_dim **<=** dim_tab**$**start_zero**[**i**])** return**(**"==="**)** **else** return**(**"・・・"**))**

dim_tab**$**end_chr **=** sapply**(**1**:**nrow**(**dim_tab**)**, **function(**i**)** **if** **(**dim_tab**$**end**[**i**]** **+** sector_dim **>=** dim_tab**$**end_zero**[**i**])** return**(**"==="**)** **else** return**(**"・・・"**))**

```
  return(dim_tab)
}
out_of_telomeres = function(dim_tab) {
  dim_tab = subset(dim_tab, (start_chr == "===" & start_zero + first_kmer > start + sector_dim) | (end_chr == "===" & start_zero + last_kmer < end - sector_dim))[,c("chr","side","chr_num","start","end","start_zero","end_zero","first_kmer","last_kmer","start_chr","end_chr")]
  out_telomeres = data.frame(chr = character(0),
                             init = numeric(0),
                             term = numeric(0),
                             lato = character(0))
  for (i in 1:nrow(dim_tab)) {
    if (dim_tab$start_chr[i] == "===") out_telomeres[nrow(out_telomeres) + 1,] = c(dim_tab$chr[i], dim_tab$start[i]-dim_tab$start_zero[i], dim_tab$first_kmer[i],paste0(dim_tab$chr[i],"_pretelomeric"))
    if (dim_tab$end_chr[i] == "===") out_telomeres[nrow(out_telomeres) + 1,] = c(dim_tab$chr[i], dim_tab$last_kmer[i], dim_tab$end[i]-dim_tab$start_zero[i],paste0(dim_tab$chr[i],"_posttelomeric"))
  }
  out_telomeres$length = abs(as.numeric(out_telomeres$init) - as.numeric(out_telomeres$term))
  out_telomeres$lengthHR = sapply(out_telomeres$length, function(o) convertir_unite(o,R = 1))
  return(out_telomeres)
}
  len_of_telomeres = function(telomeric_sectors) {
    all_sectors = Reduce(rbind.data.frame,
                         sapply(names(telomeric_sectors), function(k) {
                           cbind.data.frame(telomeric_sectors[[k]][, c('side',"chr","start_sector","end_sector","Repetitions") ],
                                            kmer = k,
                                            in_list = 1:nrow(telomeric_sectors[[k]]))
                         }, simplify = FALSE
                         ))
    cat("\n Computing lengths of telomeres \n> Assinging sectors")
    telomeric_table = Reduce(rbind.data.frame,sapply(1:nrow(all_sectors), function(x) {
      cat(progressing_bar(x,nrow(all_sectors)))
      if (x == nrow(all_sectors)) cat ("> Finishing... \n")
      K = all_sectors[x,"kmer"]
      P = unlist(kmer_this_chr[[all_sectors$kmer[x]]][all_sectors[x,"in_list"]])
      C = x
      L = length(P)
      return(data.frame(kmer = rep(K, L),
                        sect = rep(C, L),
                        posdecale = k_tab$posdecale[P])
      )
    }
    , simplify = FALSE))
    telomeric_table = merge(k_tab , telomeric_table,by = c("kmer","posdecale"))
    telomeric_table$nb_kmer = sapply(telomeric_table$sect, function(n) sum(telomeric_table$sect == n))
    telomeric_table = subset(merge(chr_dim[,c("side","start_chr","end_chr")],
                                   telomeric_table, by = "side"),
                             (start_chr == "===" | end_chr == "===") & nb_kmer > THRESHOLD_KMER_RETAIN)
    cat("> Designing output table \n")
    len_tel = as.data.frame(t(sapply(unique(telomeric_table$side), function(s) {
      this_side = subset(telomeric_table, side == s)
      L = max(this_side$pos) - min(this_side$pos)
      return(c(chr = unique(this_side$chr),
               start = min(this_side$pos),
               end = max(this_side$pos),
               length = L,
               lengthHR = convertir_unite(L,1),
               percent_chr = round(100 * L / subset(chr_zero, chr == unique(this_side$chr))$length_zero,4),
               percent_genome = round(100 * L / sum(chr_zero$length_zero),4),
               side = s
      ))
    }, simplify = TRUE)))
    len_tel$length = as.numeric(len_tel$length)
    len_tel["Total",] = c(chr = NA,
                          start = NA,
                          end = NA,
                          length = sum(len_tel$length),
                          lengthHR = convertir_unite(sum(len_tel$length),1),
                          percent_chr = NA,
                          percent_genome = 100 * sum(len_tel$length) / sum(chr_zero$length_zero),
                          side = NA
    )
    return(len_tel[order(as.numeric(len_tel$length), decreasing = TRUE),])
        }
calc_legend = function() {
  max_val = max(sector_tab$end_sector_graph)/15
  lgrth = round(log10(max_val))-1
  leg_val = round(max_val/(2*10^lgrth), 0)*2*10^lgrth
  barres = 0.1*nrow(chr_dim)/40
  pos = max(chr_dim$chr_num)+1-ESPACE_X_CHROMOSOME+barres
  min_x_pos = sector_dim + min(subset(sector_tab, chr_num == nrow(chr_dim))$start_sector_graph)
  leg = rbind.data.frame( data.frame(type = rep("segment", 3),
                                     x =  c(rep(pos-barres,2),pos),
                                     xend = c(rep(pos+barres,2),pos),
                                     y = min_x_pos + c(0,leg_val,0),
                                     yend = min_x_pos + c(0,leg_val,leg_val),
                                     ecrire = rep(NA,3)
                                     ),
                          data.frame(type = "texte",
                                     x = pos + barres,
                                     y = min_x_pos + c(leg_val/2),
                                     xend = NA, yend = NA,
                                     ecrire = convertir_unite(leg_val)
                                     )
                          )
  return(leg)
}
graphique_gigiplot = function(kmer) {
  cat(">",kmer,"\n")
  if (! exists("chr_leg")) x_place = -4:max(chr_dim$chr_num) else x_place = 1:(max(chr_dim$chr_num)+2)
  ttl = paste0(LAT_NAME,": ",
               rep(kmer, ONE_KMER == FALSE), rep(", ", iteration == 0 & ONE_KMER == FALSE),
               rep(paste0(sum(reps[,kmer])," repetitions in total"),iteration == 0),
               rep(", ", iteration > 0 & ONE_KMER == FALSE),
               rep(paste0("zoom ",iteration), iteration > 0))
  gigi_segment<-ggplot() +
    scale_fill_gradient(low = COULR_MIN, high = COULR_MAX) +
    theme_void(base_size = 5) +
    theme(
      plot.subtitle=element_blank(),
          rect = element_rect(fill = "transparent", color = "transparent"),
          plot.background = element_rect(fill = "transparent", color = "transparent"),
          panel.background = element_rect(fill = "transparent", color = "transparent"),
          plot.caption = element_text(face="plain",size=7)
          ) +
    labs(title=ttl,caption = paste0("Values inferior to ",THRESHOLD_KMER_PRINT," not printed - Sector length: ~",convertir_unite(sector_dim))) +
    xlim(min(x_place)-ESPACE_X_CHROMOSOME, max(x_place)-ESPACE_X_CHROMOSOME) +
    geom_rect(data=kmer_sector_tab[[kmer]], aes(xmin=xmin, xmax=xmax, ymin=start_sector_graph,  ymax=end_sector_graph, fill=Repetitions),color=NA) +
    geom_text(data=kmer_sector_tab[[kmer]][kmer_sector_tab[[kmer]]$"Repetitions">=THRESHOLD_KMER_PRINT,], aes(y=ylabel, x=chr_num, label=Repetitions), fontface="bold", size=1, color="black", vjust = 0.5, hjust = 0.5) +
    geom_text(data=kmer_chr_dim[[kmer]], aes(y=rep_label, x=chr_num, label=Repetitions_this_chr), size=1.5, color=COULR_MAX, vjust = -0.25) +
    geom_rect(data=kmer_chr_dim[[kmer]], aes(xmin=xmin, xmax=xmax, ymin=ymin, ymax=ymax), color="black", size = 0.2, fill=NA) +
    geom_segment(data = subset(chr_leg, type == "segment"), aes(x = x, xend = xend, y = y, yend = yend)) +
    geom_text(data = subset(chr_leg, type == "texte"), aes(x = x, y = y, label = ecrire), angle = -90, hjust = 0.5, vjust = 0, size = 1)
	if (iteration == 0) {
	  gigi_segment <- gigi_segment + theme(plot.title=element_text(hjust=0.5, face="bold", size=10)) +
	    geom_text(data = chr_dim, aes(y = slot_chr_name, x = chr_num, label = paste0(chr_num)),size=1.5, colour = "black")
	}
	if (iteration > 0) {
	  gigi_segment <- gigi_segment + theme(plot.title = element_blank()) +
	    geom_text(data = chr_dim, aes(y = slot_chr_name, x = chr_num, label = gsub("^[a-zA-Z0]+","",side)), size = 2, color = "black", angle = -90, vjust = 0.5, hjust = -0) +
	  geom_text(data = chr_dim, aes(y = slot_low_lim, x = chr_num, label = start_chr), size = 2, color = "black", hjust = 0.5) +
	  geom_text(data = chr_dim, aes(y = slot_up_lim, x = chr_num, label = end_chr), size = 2, color = "black", hjust = 0.5)
	}
  return(gigi_segment)
}
cat("\n \n \n ==== ", LAT_NAME ," ==== \n \n")
for (ONE_KMER in KMER_UNO) {
file_unico = (STAMPA_PDF | STAMPA_SVG) & ONE_KMER
if (file_unico) telomeric_segment_list <- list()
  for (iteration in 0:MAX_ITERATION) {
cat("\n\nIteration ",iteration,", Single kmer reduction : ", ONE_KMER,"\n")
	if (iteration == 0) {
	  chr_dim <- raw_chr()
	  k_tab <- raw_tab()
	} else {
	  chr_dim <- telomeric_chr()
	  telomeric_tab()
	}
	if (ONE_KMER == TRUE) k_tab = subset(k_tab, kmer == k_tab$kmer[1])
	kmer_list = unique(k_tab$kmer)
	reps <- subset(data.frame(chr = chr_dim$chr,
							   side = chr_dim$side,
							   total_nb_kmer = sapply(chr_dim$side,function(x) {
								 return(length(which(k_tab$side == x)))
								 })), total_nb_kmer != 0 )
	reps <- calc_xy(reps)
	chr_dim <- calc_xy(chr_dim)
	if (iteration == 0) {
	  chr_zero <- chr_dim[,c("chr","length","start","end","xmin","xmax")]
	  names(chr_zero) <- c("chr",paste0(grep("chr", names(chr_zero),invert = TRUE, value = TRUE),"_zero"))
	 }
	k_tab <- calc_xy(k_tab)
	sector_dim <- round(max(chr_dim$length)/MAX_SECTOR_NUMBER,0)
	if (sum(chr_dim$length<sector_dim) > 0) sector_dim <- min(chr_dim$length)
	chr_dim$sectors <- round(chr_dim$"length"/sector_dim, 0)
	esp_resp = max(chr_dim$length)*ESPACE_Y_REPS
	sector_tab <- calc_xy(calc_sectors())
	calc_kmer_sectors(k_tab$kmer)
	chr_dim <- merge(chr_dim,chr_zero,by = "chr")
	chr_dim <- calc_graph_adds(chr_dim)
	 chr_leg = calc_legend()
	if (!file_unico) telomeric_segment_list <- list()
	cat("\n Creating graphics\n")
	for (kmer in kmer_list) if (!file_unico) {
	  telomeric_segment_list[[kmer]] <- graphique_gigiplot(kmer) } else {
	    telomeric_segment_list[[as.character(iteration)]] <- graphique_gigiplot(kmer)
	    }
	if ((! STAMPA_PDF) | iteration == MAX_ITERATION) print(telomeric_segment_list[[1]])
	  if (STAMPA_PDF & (! ONE_KMER | iteration == MAX_ITERATION) ) {
    if ( ! ONE_KMER ) {
      pdf.file = paste0(OUTDIR,"/",SPECIES,"_telomeric_kmers_segments_zoom_",iteration,".pdf")
      } else {
      pdf.file = paste0(OUTDIR,"/",SPECIES,"_telomeric_kmers_segments_",kmer_list[1],".pdf")
        }
    cat(" Printing graphics ...")
    pdf(pdf.file)
    lapply(telomeric_segment_list, function(l) print)
    dev.off()
    cat(" Done.\n")
}
	if (file_unico & STAMPA_SVG) {
	    svg.file = paste0(OUTDIR,"/",SPECIES,"_zoom_",iteration,".svg")
	    cat(" Printing svg graphic ...")
	    ggsave(plot = telomeric_segment_list[[as.character(iteration)]],
	           filename = svg.file,device = "svg",width = 20, height = 10,units = "cm",dpi = "retina")
	    ggsave(plot = telomeric_segment_list[[as.character(iteration)]],
	           filename = gsub("svg","jpeg",svg.file),device = "jpeg",bg = NULL,width = 20, height = 10,units = "cm",dpi = "retina")
	    cat(" Done.\n")
	}
  if (STAMPA_TAB & !ONE_KMER) {
    if(iteration == 0) {
      stats_rep = as.data.frame(t(reps[, c(kmer_list)])) ; names(stats_rep) = reps$side
      stats_rep$Total = as.numeric(apply(stats_rep,1, sum))
      stats_rep$cumul_length = stats_rep$Total*KMER_LENGTH
      stats_rep$cumul_lengthHR = sapply(stats_rep$cumul_length, convertir_unite)
      write.table(stats_rep,paste0(OUTDIR,"/",SPECIES,"_kmer_rep_table.txt"),quote = FALSE,row.names = TRUE,col.names = TRUE,sep = '\t')
    }
    if (iteration == 1) {
      if(SPECIES == "Pinot Noir Génome T2T") {preposttel = out_of_telomeres(chr_dim)
          write.table(preposttel[,-grep("length", names(preposttel))],paste0(OUTDIR,"/",SPECIES,"_sequences_pre-post-telomeres.txt"),quote = FALSE,row.names = FALSE,col.names = FALSE,sep = '\t')
          write.table(preposttel[,-grep("lato", names(preposttel))],paste0(OUTDIR,"/",SPECIES,"_pre-post-telomeres.txt"),quote = FALSE,row.names = FALSE,col.names = TRUE,sep = '\t')}
      telomeric_lengths = len_of_telomeres(kmer_sector_tab)
      cat("\n (",SPECIES,") Total length of telomeres: ",convertir_unite(sum(as.numeric(telomeric_lengths$length[-grep("Total", rownames(telomeric_lengths))]))),"\n")
      write.table(telomeric_lengths[-grep("Total", rownames(telomeric_lengths)),c("chr","start","end","side")],paste0(OUTDIR,"/",SPECIES,"_sequences_of_telomeres.txt"),quote = FALSE,row.names = FALSE,col.names = FALSE,sep = '\t')
      write.table(telomeric_lengths,paste0(OUTDIR,"/",SPECIES,"_stats_telomeres.txt"),quote = FALSE,row.names = TRUE,col.names = TRUE,sep = '\t')
    }
    }
}
}
```
