## Supplementary File 2 for "GRASP: A PLANT TRANSFORMATION-INDEPENDENT CRISPR-BASED SYSTEM FOR AFFINITY PURIFICATION OF SPECIFIC CHROMATIN LOCI"

Spacers

RED: spacer sequences / CYAN: PAM sequences

Spacer and PAM sequences

Spacer telomeres: GGGTTTAGGGTTTAGGGTTTAGG

Spacer Gal4: AACGACTAGTTAGGCGTGTANGG

Spacer gRNA STS #1: TAGAAACGCTCAACGTGCCAAGG

Spacer gRNA STS #2: TTACATGGGCTAAAGACAAAGGG

Spacer VvActin: CAGCAGCATGAAGATCAAGGTGG

Primers

>Tel_SHRT_Fw

GGGTTTAGGGTTTAGGGTTTAGGCAATCTCTTAGTCGACT

>Gal_SHRT_Fw

AACGACTAGTTAGGCGTGTANGGCAATCTCTTAGTCGACT

>SHRT_Rv

CTCCGTTTTACCTGTGGAATCG

>LONG_Fw

CAGGAAACAGCTATGAC

>Tel_LONG_Rv

CCTAAACCCTAAACCCTAAACCCTTATATTCCCCAGAACATCAGGTTAATGG

>Gal_LONG_Rv

CCNTACACGCCTAACTAGTCGTTTTATATTCCCCAGAACATCAGGTTAATGG

>ampl_fram_Fw

TGTGCCTGTCTCCGTTATCG

>ampl_fram_Rv

GCAGCCTGACAGACAAATGAG

>gR-R

CGGAGGAAAATTCCATCCAC

>gRT-gRNA_STS_1_Fw

TAGAAACGCTCAACGTGCCAGTTTTAGAGCTAGAAAT

>gRT-gRNA_STS_2_Fw

TTACATGGGCTAAAGACAAAGTTTTAGAGCTAGAAAT

>gRT-gRNA_VvActin_Fw

CAGCAGCATGAAGATCAAGGGTTTTAGAGCTAGAAAT

>T7-gRNA_STS_1_Fw

CCTCTAATACGACTCACTATAGGTAGAAACGCTCAACGTGCCA

>T7-gRNA_STS_2_Fw

CCTCTAATACGACTCACTATAGGTTACATGGGCTAAAGACAAA

>T7-gRNA_VvActin_Fw

CCTCTAATACGACTCACTATAGGCAGCAGCATGAAGATCAAGG

>gRNA_STS_1_SHRT_Fw

TAGAAACGCTCAACGTGCCAAGGCAATCTCTTAGTCGACT

>gRNA_STS_2_SHRT_Fw

TTACATGGGCTAAAGACAAAGGGCAATCTCTTAGTCGACT

>gRNA_VvActin_SHRT_Fw

CAGCAGCATGAAGATCAAGGTGGCAATCTCTTAGTCGACT

>gRNA_STS_1_LONG_Rv

CCTTGGCACGTTGAGCGTTTCTATTATATTCCCCAGAACATCAGGTTAATGG

>gRNA_STS_2_LONG_Rv

CCCTTTGTCTTTAGCCCATGTAATTATATTCCCCAGAACATCAGGTTAATGG

>gRNA_VvActin_LONG_Rv

CCACCTTGATCTTCATGCTGCTGTTATATTCCCCAGAACATCAGGTTAATGG

Fragments

>Tel_Long_region

CAGGAAACAGCTATGACCATGATTACGCCAAGCTATTTAGGTGACACTATAGAATACTCAAGCTATGCATCAAGCTCAATGGGTCTAGTCTGTAGATACCCATCACACTGGCGACCGCTCGAACATCAGTTTAAGGTTTACACCTATAAAAGAGAGAGCCGTTATCGTCTGTTTGTGGATGTACAGAGTGATATTATTGACACGCCGGGGCGACGGATGGTGATCCCCCTGGCCAGTGCACGTCTGCTGTCAGATAAAGTCTCCCGTGAACTTTACCCGGTGGTGCATATCGGGGATGAAAGCTGGCGCATGATGACCACCGATATGGCCAGTGTGCCTGTCTCCGTTATCGGGGAAGAAGTGGCTGATCTCAGCCACCGCGAAAATGACATCAAAAACGCCATTAACCTGATGTTCTGGGGAATATAAGGGTTTAGGGTTTAGGGTTTAGG

>Gal_Long_region

GACGAAACAGCTATGACCATGATTACGCCAAGCTATTTAGGTGACACTATAGAATACTCAAGCTATGCATCAAGCTCAATGGGTCTAGTCTGTAGATACCCATCACACTGGCGACCGCTCGAACATCAGTTTAAGGTTTACACCTATAAAAGAGAGAGCCGTTATCGTCTGTTTGTGGATGTACAGAGTGATATTATTGACACGCCGGGGCGACGGATGGTGATCCCCCTGGCCAGTGCACGTCTGCTGTCAGATAAAGTCTCCCGTGAACTTTACCCGGTGGTGCATATCGGGGATGAAAGCTGGCGCATGATGACCACCGATATGGCCAGTGTGCCTGTCTCCGTTATCGGGGAAGAAGTGGCTGATCTCAGCCACCGCGAAAATGACATCAAAAACGCCATTAACCTGATGTTCTGGGGAATATAAAACGACTAGTTAGGCGTGTANGG

>Tel_Short_region

GGGTTTAGGGTTTAGGGTTTAGGCAATCTCTTAGTCGACTCTACCAATATATAAACAGAGCTACTATTTTCAACTGAACGATGTGGGATGTTTTTGACATCGTTTACAGCTTAAACGGGCTCGTTTATTAAAAGCCCTCTTCTTTCGATCCATCAACATTATTGGCCTTAAGTAAAACAAGCCTCTTTATGATTGAGAAAACGCCGTCGTTGGTGCACCAGAGAGAAGAAACAAAACAGCTAGCGTTAGATAAACTTTAAAATTGTTTGCATCCTCATTTGTCTGTCAGGCTGCAGTAGTTTGGATTAAGAACGAAACTAACTGTAAGAGATCTCTGATATTTTCCTTTGCTGCCGATTCCACAGGTAAAACGGAG

>Gal_Short_region

AACGACTAGTTAGGCGTGTANGGCAATCTCTTAGTCGACTCTACCAATATATAAACAGAGCTACTATTTTCAACTGAACGATGTGGGATGTTTTTGACATCGTTTACAGCTTAAACGGGCTCGTTTATTAAAAGCCCTCTTCTTTCGATCCATCAACATTATTGGCCTTAAGTAAAACAAGCCTCTTTATGATTGAGAAAACGCCGTCGTTGGTGCACCAGAGAGAAGAAACAAAACAGCTAGCGTTAGATAAACTTTAAAATTGTTTGCATCCTCATTTGTCTGTCAGGCTGCAGTAGTTTGGATTAAGAACGAAACTAACTGTAAGAGATCTCTGATATTTTCCTTTGCTGCCGATTCCACAGGTAAAACGGAG

>Tel_Complete_fragment

CAGGAAACAGCTATGACCATGATTACGCCAAGCTATTTAGGTGACACTATAGAATACTCAAGCTATGCATCAAGCTCAATGGGTCTAGTCTGTAGATACCCATCACACTGGCGACCGCTCGAACATCAGTTTAAGGTTTACACCTATAAAAGAGAGAGCCGTTATCGTCTGTTTGTGGATGTACAGAGTGATATTATTGACACGCCGGGGCGACGGATGGTGATCCCCCTGGCCAGTGCACGTCTGCTGTCAGATAAAGTCTCCCGTGAACTTTACCCGGTGGTGCATATCGGGGATGAAAGCTGGCGCATGATGACCACCGATATGGCCAGTGTGCCTGTCTCCGTTATCGGGGAAGAAGTGGCTGATCTCAGCCACCGCGAAAATGACATCAAAAACGCCATTAACCTGATGTTCTGGGGAATATAAGGGTTTAGGGTTTAGGGTTTAGGCAATCTCTTAGTCGACTCTACCAATATATAAACAGAGCTACTATTTTCAACTGAACGATGTGGGATGTTTTTGACATCGTTTACAGCTTAAACGGGCTCGTTTATTAAAAGCCCTCTTCTTTCGATCCATCAACATTATTGGCCTTAAGTAAAACAAGCCTCTTTATGATTGAGAAAACGCCGTCGTTGGTGCACCAGAGAGAAGAAACAAAACAGCTAGCGTTAGATAAACTTTAAAATTGTTTGCATCCTCATTTGTCTGTCAGGCTGCAGTAGTTTGGATTAAGAACGAAACTAACTGTAAGAGATCTCTGATATTTTCCTTTGCTGCCGATTCCACAGGTAAAACGGAG

>Gal_Complete_fragment

GACGAAACAGCTATGACCATGATTACGCCAAGCTATTTAGGTGACACTATAGAATACTCAAGCTATGCATCAAGCTCAATGGGTCTAGTCTGTAGATACCCATCACACTGGCGACCGCTCGAACATCAGTTTAAGGTTTACACCTATAAAAGAGAGAGCCGTTATCGTCTGTTTGTGGATGTACAGAGTGATATTATTGACACGCCGGGGCGACGGATGGTGATCCCCCTGGCCAGTGCACGTCTGCTGTCAGATAAAGTCTCCCGTGAACTTTACCCGGTGGTGCATATCGGGGATGAAAGCTGGCGCATGATGACCACCGATATGGCCAGTGTGCCTGTCTCCGTTATCGGGGAAGAAGTGGCTGATCTCAGCCACCGCGAAAATGACATCAAAAACGCCATTAACCTGATGTTCTGGGGAATATAAAACGACTAGTTAGGCGTGTANGGCAATCTCTTAGTCGACTCTACCAATATATAAACAGAGCTACTATTTTCAACTGAACGATGTGGGATGTTTTTGACATCGTTTACAGCTTAAACGGGCTCGTTTATTAAAAGCCCTCTTCTTTCGATCCATCAACATTATTGGCCTTAAGTAAAACAAGCCTCTTTATGATTGAGAAAACGCCGTCGTTGGTGCACCAGAGAGAAGAAACAAAACAGCTAGCGTTAGATAAACTTTAAAATTGTTTGCATCCTCATTTGTCTGTCAGGCTGCAGTAGTTTGGATTAAGAACGAAACTAACTGTAAGAGATCTCTGATATTTTCCTTTGCTGCCGATTCCACAGGTAAAACGGAG

>Tel_Shortened_fragment

TGTGCCTGTCTCCGTTATCGGGGAAGAAGTGGCTGATCTCAGCCACCGCGAAAATGACATCAAAAACGCCATTAACCTGATGTTCTGGGGAATATAAGGGTTTAGGGTTTAGGGTTTAGGCAATCTCTTAGTCGACTCTACCAATATATAAACAGAGCTACTATTTTCAACTGAACGATGTGGGATGTTTTTGACATCGTTTACAGCTTAAACGGGCTCGTTTATTAAAAGCCCTCTTCTTTCGATCCATCAACATTATTGGCCTTAAGTAAAACAAGCCTCTTTATGATTGAGAAAACGCCGTCGTTGGTGCACCAGAGAGAAGAAACAAAACAGCTAGCGTTAGATAAACTTTAAAATTGTTTGCATCCTCATTTGTCTGTCAGGCTGC
