## Supplementary Figures for "GRASP: A PLANT TRANSFORMATION-INDEPENDENT CRISPR-BASED SYSTEM FOR AFFINITY PURIFICATION OF SPECIFIC CHROMATIN LOCI"

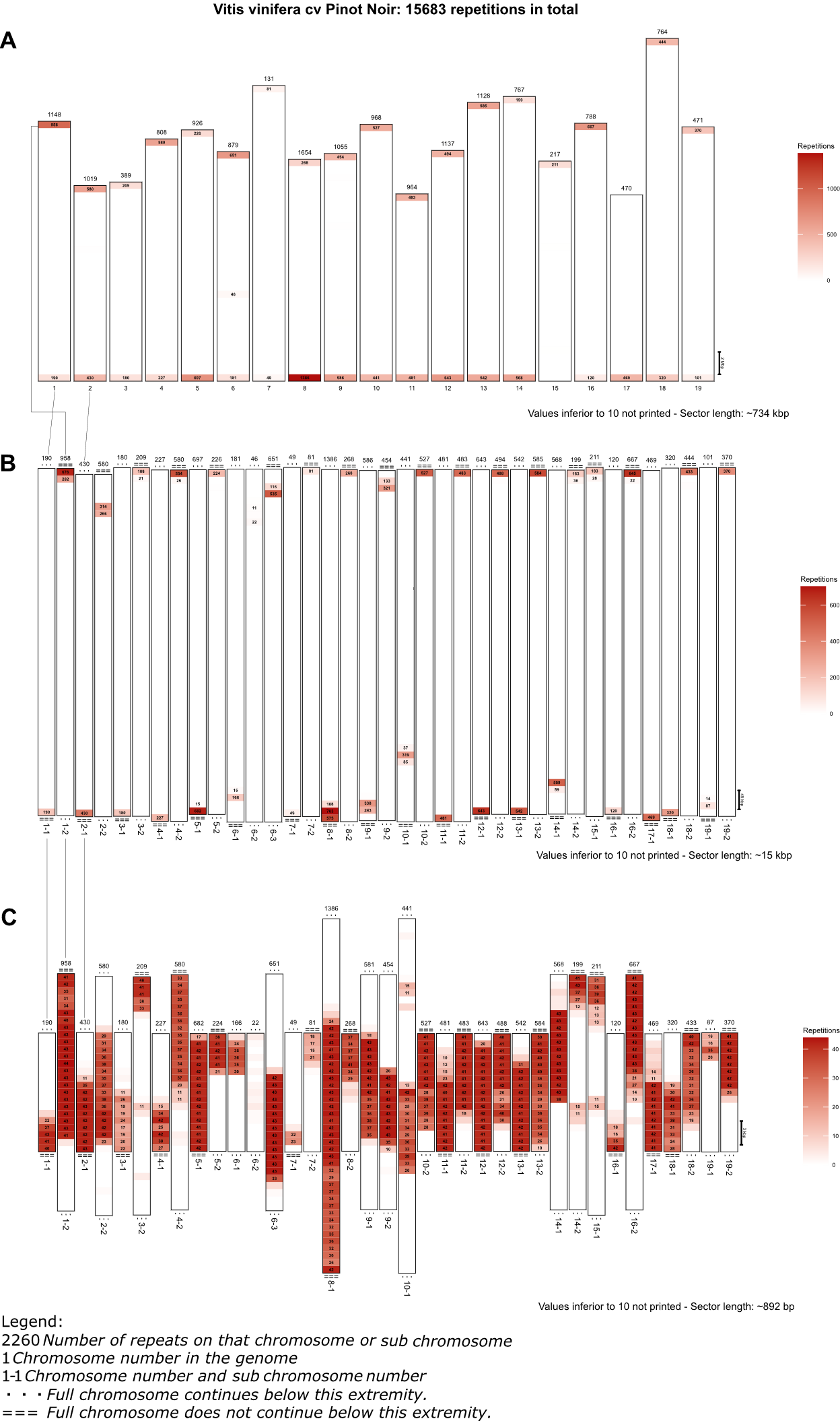

**Figure S1.** Abundance of telomeres in the T2T genomes of Vitis vinifera cv Pinot Noir. Repeats of the 21-kmer (here AAACCCTAAACCCTAAACCCT) were blasted against the genomes of the Pinot Noir cultivar, and the resulting positions placed on the chromosomes are divided into sectors (**A**). A zoom was subsequently applied to the sectors containing at least 20 repetitions (**B**). A further zoom was applied again to the sectors containing 20 or more repetitions of the kmer (**C**).

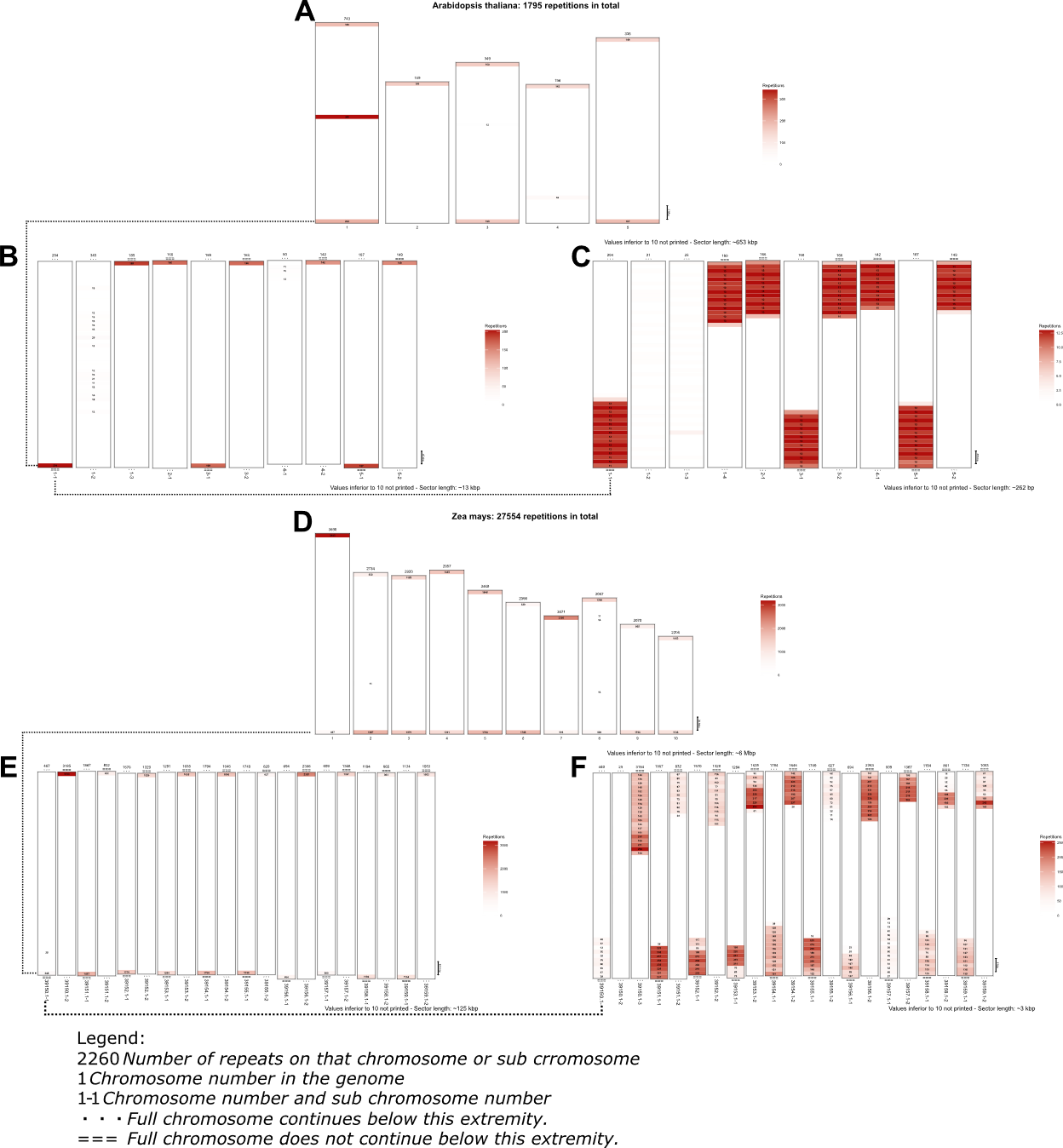

**Figure S2.** Abundance of telomeres in the T2T genomes of two non-*Vitis* species. Repeats of the 21-kmer (here AAACCCTAAACCCTAAACCCT) where blasted against the genomes of *Arabidopsis thaliana* (**A-C**) and *Zea mays* (**D-F**), and the resulting positions placed on the chromosomes are divided into sectors (**A,D**). A zoom was subsequently applied to the sectors containing at least 20 repetitions (**B,E**). A further zoom was applied again to the sectors containing 20 or more repetitions of the kmer.

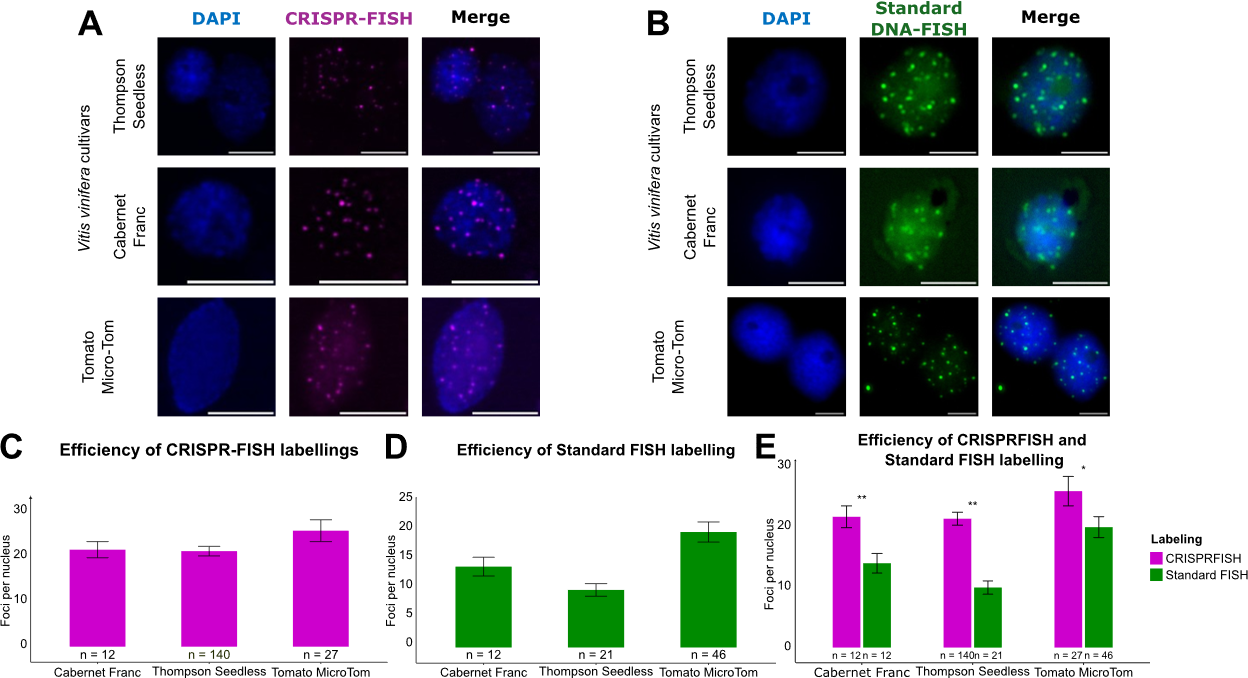

**Figure S3.** CRISPR-FISH and Standard FISH on grapevine and tomato nuclei. We performed CRISPR-FISH (**A**) and standard DNA FISH (**B**) on nuclei of grapevine cv. Thompson Seedless and Cabernet Franc and Tomato MicroTom. We used an ImageJ macro to count the foci numbers for CRISPR–FISH (**C**) and standard-FISH (**D**) on our images. (**E**) Comparison between the two labeling methods. For statistical analysis, we performed unpaired Student’s t tests. * p < 0.05, ** p < 0,01. The error bars represent the standard error. Scale bar: 8 µm.

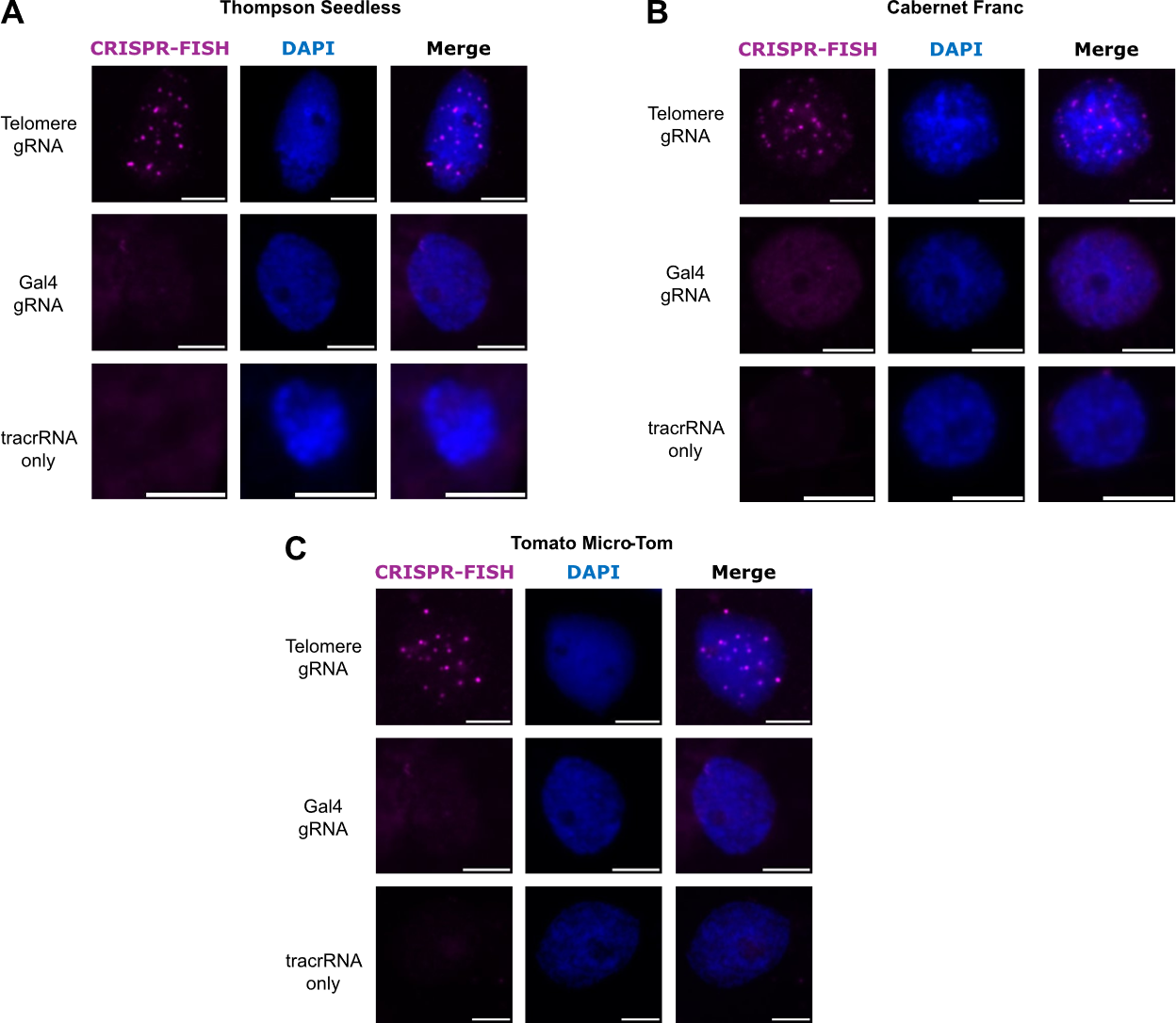

**Figure S4.** CRISPR–FISH on grapevine and tomato nuclei including non-targeting controls. We performed CRISPR-FISH on nuclei of grapevine Thompson Seedless, Cabernet Franc and Tomato Micro-Tom using the full ATTO-labelled gRNA targeting either telomeres or bacterial Gal4 gene, or only the ATTO-labeled tracrRNA. Scale-bar: 8µm.

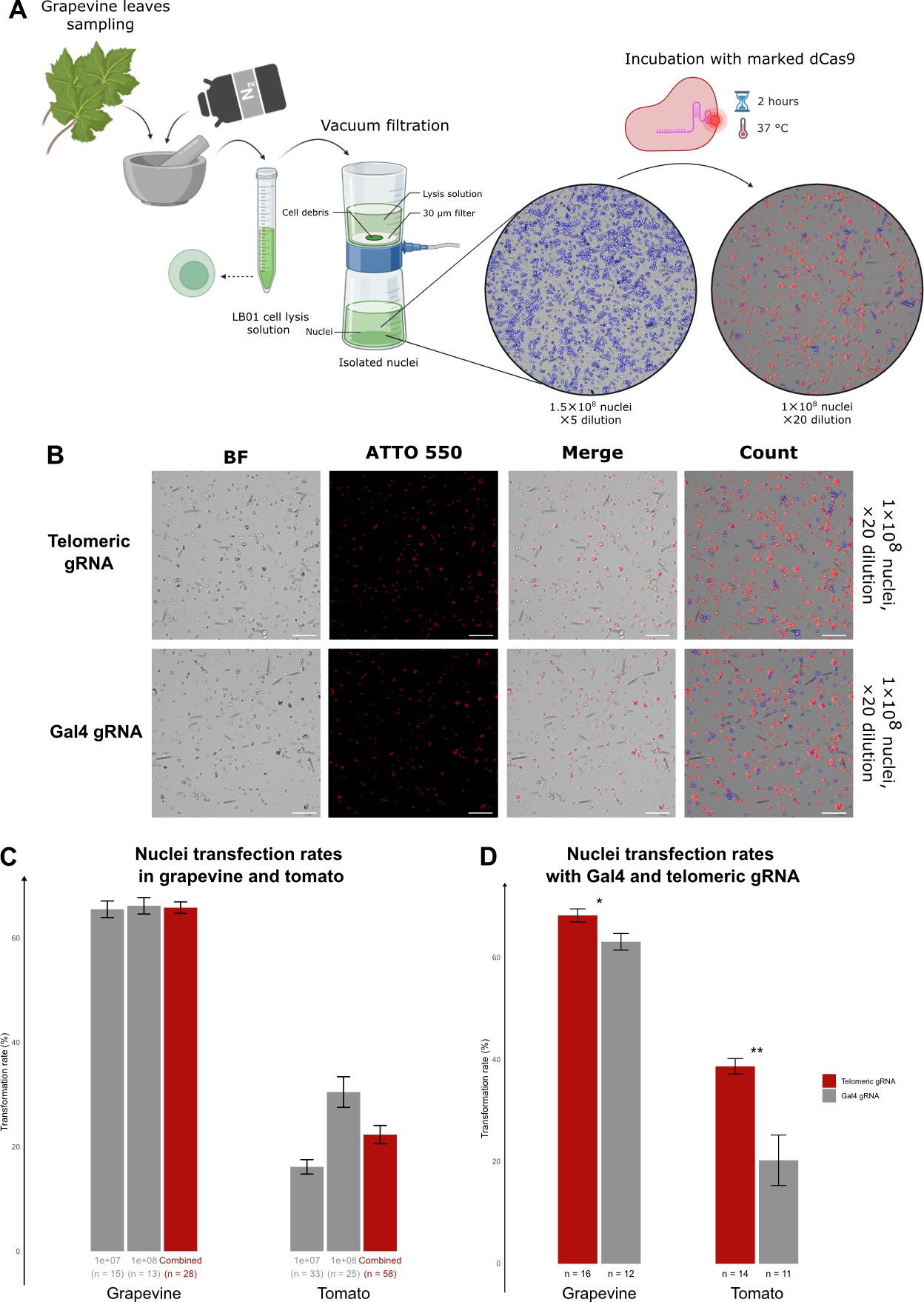

**Figure S5.** Nuclei isolation from grapevine leaves, transformation and counting. (**A**) Schematical description of the experiment: grapevine nuclei were isolated and counted using a CellDrop cell counter. Then, 10^7^ or 10^8^ nuclei per sample were incubated with dCas9 loaded with a gRNA against telomeres carrying an ATTO550 fluorophore, and the transfection efficiency was evaluated with the same cell counter. (**B**) Images of grapevine nuclei transfected with dCas9 and ATTO-gRNA against telomeres or Gal4 obtained from the cell counter. “BF”, “ATTO550” and “Count”-type images were produced by the instrument, whereas “Merge”-type images were produced by merging BF and ATTO550 channels manually. The number of nuclei indicates the total number of nuclei counted in the whole sample before incubation with dCas9. “Dilution” is the dilution applied to the sample before counting (i.e., for a 20-fold dilution, 1 µL of sample was diluted into 19 µL of water, and 10 µL of such dilution was loaded on the cell counter). (**C**) Transfection rates in tomato and grapevine with gRNA against telomeres as a function of the number of nuclei (**D**) Transfection rates in tomato (samples of (**C**) containing 10^8^ nuclei) and grapevine (samples of (**C**) containing 10^7^ and 10^8^ nuclei) as a function of the gRNA used. * p < 0,05 ** p < 0,001, Student’s t test, number of measures: indicated under the corresponding boxplot.

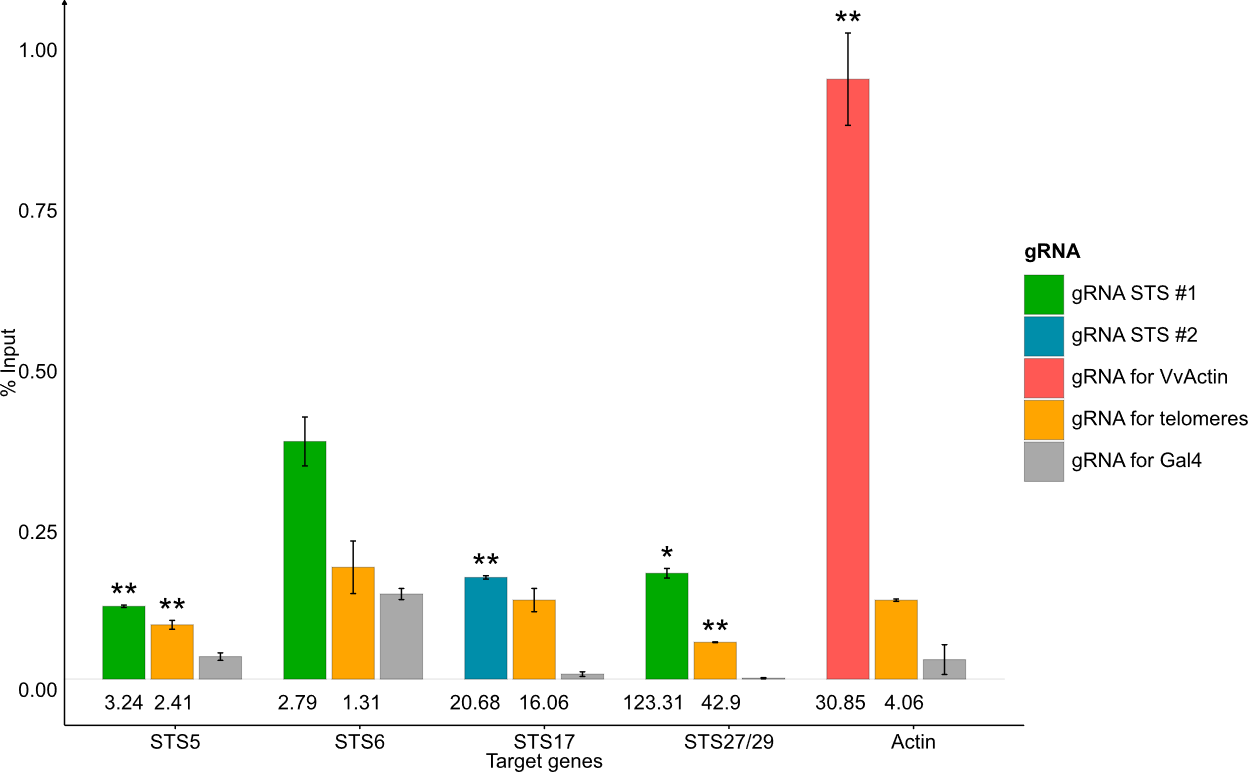

**Figure S6.** Precipitation of lesser-repeated spacers with telomeric gRNA as secondary control. Precipitation was performed on nuclei of grapevine cv Thompson Seedless, using 10^8^ nuclei for each transfection and one of each following gRNA: gRNA STS #1, gRNA STS #2, gRNA for VvActin gRNA for telomeres and gRNA for Gal4. qPCR on target genes for which the gRNAs are specific (gRNA STS #1: STS5, STS6, STS27/29 ; gRNA STS #2: STS17; gRNA for VvActin: Actin) was performed in the precipitated and in the input samples. Plus, qPCR of all these target genes was performed in the precipitated and input sample with telomeric and Gal4 gRNA. Mean CTs in precipitated were then normalized with the counterparts in input, and the percentage of input represented by the precipitated was plotted for each gen in the cases when the gRNA used for precipitation is specific of that gene (Specific, colored) or when the used gRNA is a negative control telomeres (unspecific, orange) or Gal4 gRNA (no target, gray). Values under the bars indicate the %input fold-change between the considered precipitation and Gal4 precipitation. Significativity of differences was evaluated with Student t-test ($*$ p < 0.05, $**$ p < 0,01).
